# Localized estimation of event-related neural source activity from simultaneous MEG-EEG with a recurrent neural network

**DOI:** 10.1101/2024.03.01.582924

**Authors:** Jamie A. O’Reilly, Judy D. Zhu, Paul F. Sowman

## Abstract

Estimating intracranial current sources underlying the electromagnetic signals observed from extracranial sensors is a perennial challenge in non-invasive neuroimaging. Established solutions to this inverse problem treat time samples independently without considering the temporal dynamics of event-related brain processes. This paper describes current source estimation from simultaneously recorded magneto- and electro-encephalography (MEEG) using a recurrent neural network (RNN) that learns sequential relationships from neural data. The RNN was trained in two phases: (1) pre-training and (2) transfer learning with L1 regularization applied to the source estimation layer. Performance of using scaled labels derived from MEEG, magnetoencephalography (MEG), or electroencephalography (EEG) were compared, as were results from volumetric source space with free dipole orientation and surface source space with fixed dipole orientation. Exact low-resolution electromagnetic tomography (eLORETA) and mixed-norm L1/L2 (MxNE) source estimation methods were also applied to these data for comparison with the RNN method. The RNN approach outperformed other methods in terms of output signal-to-noise ratio, correlation and mean-squared error metrics evaluated against ground-truth event-related field (ERF) and event-related potential (ERP) waveforms. Using MEEG labels with fixed-orientation surface sources produced the most consistent estimates. To estimate sources of ERF and ERP waveforms, the RNN generates temporal dynamics within its internal computational units, driven by sequential structure in neural data used as training labels. It thus provides a data-driven model of computational transformations from psychophysiological events into corresponding event-related neural signals, which is unique among MEG and EEG source reconstruction solutions.

## 1. Introduction

A goal of non-invasive neuroimaging is to localize sources of brain activity observed from outside the head. Localizing sources of magnetoencephalography (MEG) and electroencephalography (EEG) signals is known as the inverse problem, which is ill-posed because there are theoretically infinite compositions of source signals that can explain observed MEG or EEG signals (Nunez & Srinivasan, 2009). Due to the non-uniqueness of inverse problem solutions, constraints based on biophysics or numerical methods are employed to reduce the search space of possible solutions. The forward problem, estimating MEG and EEG signal amplitudes from an observed brain source signal, is comparatively well posed and can be estimated with knowledge of the physical conduction of electromagnetic phenomena through biological tissues (Nunez & Srinivasan, 2009). However, brain source signals are generally inaccessible, making solutions to the inverse problem desirable for estimating brain source signals that account for observed MEG and EEG signals. A *lead field* matrix of coefficients can be determined from a forward conduction model, which describes the contributions of currents in the brain to MEG and EEG sensors if head and sensor geometry are known (Hämäläinen & Ilmoniemi, 1994). The lead field matrix provides a fundamental constraint for inverse solutions. Utilizing simultaneously recorded MEG and EEG (i.e., MEEG) constrains the space of solutions to the inverse problem, which is considered superior to using either MEG or EEG on their own (Dale et al., 1993; Luck, 2014; Nunez & Srinivasan, 2009).

Inverse solutions are optimization problems. Equivalent current dipole fitting algorithms aim to determine the optimal location, orientation and moment of one or more dipoles that can account for observed data (Ahlfors & Hämäläinen, 2012). Drawbacks of dipole fitting methods are dependence of solutions on initial conditions and assuming a low number of active dipoles. The latter drawback of dipole fitting methods can be overcome by distributed source space methods, which attempt to estimate signals at many locations (referred to as vertices) in parallel (Ahlfors & Hämäläinen, 2012). Pseudoinverse matrices can be computed from the lead field matrix using methods from linear algebra, such as minimum-norm estimates (MNE; Hämäläinen & Ilmoniemi, 1994) and more advanced variants such as exact low-resolution electromagnetic tomography (eLORETA; Pascual-Marqui et al., 2011), that include different constraints (e.g., L2 norm in conventional MNE) to solve the inverse problem. For instance, the L1/L2 mixed-norm estimates (MxNE) method includes an L1 norm penalty to obtain a sparse solution to the inverse problem (Gramfort et al., 2012). However, the rationale and reliability of these numerical constraints for determining the true distribution of brain currents remains open for debate (Luck, 2014).

Methods of MEG and EEG source localization have also arrived from Bayesian probability theory (Das et al., 2020; David et al., 2006; Henson et al., 2010; Henson, Mouchlianitis, et al., 2009; Wipf & Nagarajan, 2009). These have incorporated different priors to constrain inverse solutions, such as preconceived network topologies and interactions (Kiebel et al., 2006), constraints derived from function magnetic resonance imaging data (Henson et al., 2010), and deconvolving signals using a bank of linear “neuro-current response functions” (Das et al., 2020). Several methods have also emerged from recent developments in deep learning with artificial neural networks, reportedly having equivalent or superior performance to conventional linear inverse solutions (Hecker et al., 2021; Liang et al., 2023; Pantazis & Adler, 2021; Sun et al., 2022). However, these methods do not extract temporal structure from neural data because they are fitted to an individual or small number of time points, which limits the extent to which resulting solutions can be interpreted.

Recurrent neural network (RNN) models are designed to learn sequential relationships from data, potentially making them more useful for analysing neural time series than other artificial neural networks. Simple RNNs have been used to model event-related neural signals from mice exposed to auditory stimulation (O’Reilly, 2022a; O’Reilly, Angsuwatanakul, et al., 2022) and humans exposed to auditory and visual stimuli (O’Reilly, 2022b; O’Reilly, Wehrman, et al., 2023). A long short-term memory (LSTM; Hochreiter & Schmidhuber, 1997) type of RNN has been used to refine MEG and EEG source estimates obtained by applying MNE to individual time samples, highlighting the importance of temporal context among neural signals for improving source reconstruction (Dinh et al., 2021). Simple RNN models have also been developed for estimating localized source signals from MEG (O’Reilly, Zhu, et al., 2023) and EEG (Srivastava et al., 2023) by fixing the weights of a feed-forward output layer equal to the lead field matrix. By training the RNN in two steps: (1) pretraining and (2) transfer learning with L1 regularization, estimated source signals are derived from penultimate layer hidden unit activations (O’Reilly, Zhu, et al., 2023).

In this study, the RNN method of neural current source localization was developed for and applied to simultaneously recorded MEEG signals. Estimated source signals were obtained from MEEG, MEG, and EEG data, and two linear inverse solutions (eLORETA and MxNE) were applied to the same data for comparison. Furthermore, volumetric source space with free dipole orientation and surface source space with fixed surface-normal orientation were evaluated. The results of this study will inform future developments of RNN models of event-related neural signal processing.

## 2. Materials and methods

### 2.1. MEEG recordings

MEG and EEG were simultaneously recorded at the KIT-Macquarie Brain Research Laboratory in Sydney, Australia, using equipment described in a recent article (Zhu et al., 2022). During the recording, subjects listened to “ba” and “da” syllable sounds played in a random sequence. This experiment was part of a larger study approved by the Human Research Ethics Committee at Macquarie University. Before recording signals, an EEG cap of appropriate size was fitted to the subject’s head, and five head position marker coils were attached onto the cap (to locate the head position once inside the MEG sensor array). Subjects’ head shape information was obtained by 3D head scanning using the Structure Sensor accessory (Occipital, Inc., Boulder, CO) on an iPad, and the positions of fiducials and marker coils were marked manually. The sequence of sounds in the experiment was generated using custom code in Matlab (version R2017b; The MathWorks Inc., Natick, MA, USA) and delivered binaurally via pneumatic earphones (Etymotic ER30; Etymotic Research, Inc., Elk Grove Village, IL, USA).

The MEG data were recorded using a system with 160 axial gradiometers (Model 60R-N2; Kanazawa Institute of Technology, Kanazawa, Japan) located inside a magnetically shielded room (Fujihara Co. Ltd., Tokyo, Japan). Signals were recorded from three additional reference sensors, which were used for post-hoc denoising of MEG signals using time-shift regression (de Cheveigné & Simon, 2007). Audio signals recorded synchronously with MEG were used to obtain accurate onset timing for the auditory stimuli. The subject’s head position was measured before and after the experimental task and it deviated by less than 5 mm. Signals were acquired from the MEG system at 1000 samples per second with online band-pass filtering from 0.03 to 200 Hz.

The EEG data were recorded using an MEG-compatible 64-channel system (BrainAmp; Brain Products, Germany). The EEG cap (BrainCap-MEG; EasyCap, Germany) contained 61 EEG channels, placed according to the 10-20 system (Fp1, AF3, AF7, Fz, F1, F3, F5, F7, FC1, FC3, FC5, FT7, Cz, C1, C3, C5, T7, CP1, CP3, CP5, TP7, TP9, Pz, P1, P3, P5, P7, PO3, PO7, Oz, O1, Fpz, Fp2, AF4, AF8, F2, F4, F6, F8, FC2, FC4, FC6, FT8, C2, C4, C6, T8, CPz, CP2, CP4, CP6, TP8, TP10, P2, P4, P6, P8, POz, PO4, PO8, and O2), plus EOG, ECG, and a reference channel at FCz. Data were re-referenced offline to the average montage. An affine transform aligned default EEG channel locations with fiducials (nasion and bilateral preauricular areas) and head landmarks (vertex and occipital point) obtained from head digitization data. Channel location and head digitization alignments are plotted in Figure S1. Fiducials and head shape points were also co-registered with a template head MRI to compute forward conduction models for both MEG and EEG, as described in section 2.2.

Independent component analysis (ICA) was used to correct artifacts related to ocular, cardiological, and non-biological sources from both signal modalities (Jung et al., 2000); three components were removed from MEG, and six components were removed from EEG. EEG and MEG were band-pass filtered from 1 to 40 Hz, and resampled to 100 Hz, before extracting epochs from −0.1 to 0.4 s relative to stimulus onset on each trial. Baseline correction was applied using the pre-stimulus interval. The resulting MEEG data matrix comprised 99 “ba” epochs and 98 “da” epochs, ***Y**_MEEG_* ∈ ℝ ^197×51×221^ (total 197 epochs, 51 time-samples, and 221 channels); MEG and EEG submatrices were extracted from this, where ***Y**_MEG_* ∈ ℝ^197×51×160^ and ***Y*** ∈ ℝ^197×51×61^. These data matrices were scaled and used as labels for training RNN models.

### 2.2. Source spaces and forward models

Volume and surface source spaces were used, and the results from both were compared to assess the reproducibility of source signals estimated by either approach using the RNN method. The volume source space had 30.0 mm grid spacing with 3.0 mm minimum distance from the inner skull; producing 50 vertices, each with three components for x, y, and z directions. The surface source space had “ico3” icosahedron decimation, producing 66 vertices in each hemisphere, with orientation fixed to the cortical surface normal. While not matched exactly, these two source spaces provided similar degrees of freedom for fitting observed signals; 150 for the volume space and 132 for the surface space. More degrees of freedom might be assumed to yield better-fitting solutions to the data if that were the only difference between the two models. The average minimum distance between each vertex and its nearest neighbour was 30.0 mm for the volume source space and 15.3 mm for the surface source space.

A three-layer boundary element method (BEM) conduction model was computed from the co-registered template head, with brain, skull, and scalp conductance of 0.3, 0.006, and 0.3 S/m, respectively. Forward solutions for volume and surface source spaces were derived from this conduction model, with lead field coefficients calculated for 160 MEG and 61 EEG channels. The combined volume source space lead field matrix can be denoted 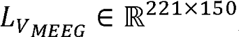, with 221 channels and 50 × 3 source signal components (x, y, and z directions for each vertex). Similarly, the combined surface source space, 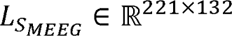, accounted for 66 sources on each hemisphere with fixed surface-normal orientation. Standardized (Z-scored) lead field coefficient distributions for MEG and EEG channels in volume and surface source spaces are plotted in Figure S2.

### 2.3. RNN inputs and labels

Figure 1 illustrates how an RNN was used to model ERF and ERP waveforms derived from simultaneously recorded MEEG. Inputs consisted of two signal channels: one representing “ba” and another representing “da” events; ***x*** ∈ ℝ^51×2^. A unit step-pulse set high from 0.0 to 0.075 s on the respective channel signalled that event. Each input was paired with a single epoch of MEEG data recorded in response to the associated event (i.e. ***y*** ∈ ℝ^51×221^), which provided labels for supervised learning. All of the single epochs in the data matrix were paired with one of two input representations, producing an input matrix, ***X*** ∈ ℝ^197×51×221^. These pairings effected a one-to-many mapping of input (encoding event type) to output label (single-trials of brain signals) time sequences, where the RNN models the function: ***X*** → ***Y***. As an artificial neural network minimizes mean-squared error loss with a one-to-many relation between its inputs and labels, its output will converge towards the average of those labels associated with each unique event. This behaviour is desirable in the context of modelling averaged event-related neural signals. The same approach was used to model waveforms from MEG-only and EEG-only by setting the model output weights (***W**_out_*) using the appropriate lead field matrix and corresponding single-modality data as labels, described in section 2.5.

**Figure 1.**
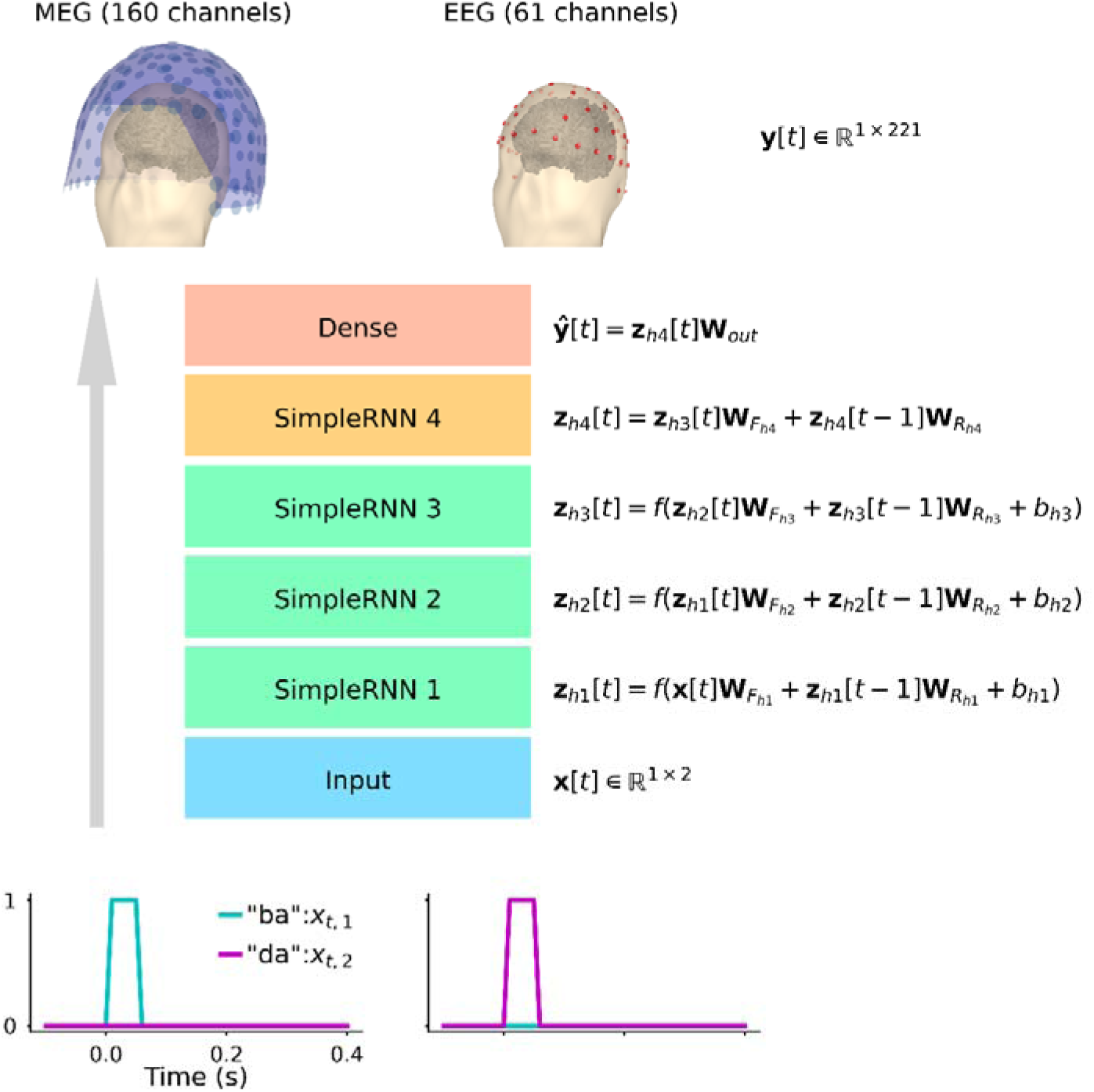
Model architecture and training scheme for using an RNN to estimate sources of MEEG data. Two sets of step pulse functions (bottom) representing “ba” (left) and “da” (right) events were input to the RNN. These inputs were transformed through sequential recurrent layers (SimpleRNN 1-4), the first three with rectified linear unit activation function, the fourth with linear activation, and then a single feed-forward (Dense) layer with linear activation. The fourth SimpleRNN layer was trained without a bias and had L1-norm activity regularization when the model was fine-tuned. The layer naming convention adopted here is consistent with the TensorFlow API. Labels for supervised learning were obtained from simultaneously recorded MEG (top-left) and EEG (top-right). The final layer of this model effectively implements the forward solution. Thus, outputs from the penultimate layer (**z**_h4_) can be interpreted as source signals. Mathematical notations illustrate the data entities and computations at each stage in the modelling process for an individual time sample.

### 2.4. RNN architecture and training parameters

As with previous RNN models of event-related neural signals (O’Reilly, 2022b; O’Reilly, Angsuwatanakul, et al., 2022; O’Reilly, Wehrman, et al., 2022; O’Reilly, Zhu, et al., 2023), the architecture consisted of an input, four hidden layers, and an output layer. Time-domain signals were passed to the input layer, transformed sequentially through four simple recurrent layers, and then fed through a feed-forward output layer. Mathematical objects and operations at each layer are annotated on the diagram in Figure 1. The first three recurrent layers had rectified linear unit activation function. The fourth recurrent layer, which generates estimated source signals, had linear activation. The output layer also had linear activation and effectively implemented the forward computation: *signals = sources × lead field*, formulated as 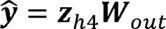, where 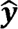 and ***W**_out_* were obtained by scaling data labels and corresponding lead field coefficients, respectively, as explained in section 2.5.

The backpropagation through time algorithm (Werbos, 1990) with adaptive moment estimation optimizer (Kingma & Ba, 2015) and mean-squared error loss were used to train the RNN, all using default hyperparameter settings in TensorFlow (Abadi et al., 2016). As described previously (O’Reilly, Zhu, et al., 2023; Srivastava et al., 2023), to derive source estimates, the RNN was pre-trained to fit labels without additional constraints (training step 1) before discarding the bias parameter and introducing L1-norm activity regularization to the fourth recurrent layer (training step 2). This L1-norm penalty was weighted by 10^−4^. In both training steps the maximum number of training epochs/iterations was set to 5000. However, training ceased if the loss failed to improve for 50 consecutive epochs; all models stopped before reaching the maximum (Figure S3). Batch size of 197 was selected to include all training instances. Due to stochastic variation in network initialization and optimization algorithms, models were trained five times using different random seeds (integers 0 to 4).

### 2.5. Scaling data and lead field

To avoid the vanishing gradients problem (Bengio et al., 1994), MEG and EEG data were scaled before being used as labels. Magnetic fields measured from the brain are commonly reported on the order of femtoteslas (fT = ×10^−15^ T). In contrast, potential differences due to electric fields of the brain are observed on the order of microvolts (μV = ×10^−6^ V). Different orders of magnitude reflect different signal conduction and acquisition physics for magnetic and electric fields, although both signals arise from biophysical processes operating on the same scale. That is to say, if synchronous brain processes are considered to generate an electric dipole that can be measured from outside the head by EEG and/or MEG, the same dipole current source density produces EEG signals in the μV range and MEG signals in the fT range. Therefore, when scaling MEG and EEG labels for supervised learning, their corresponding lead field coefficients were also scaled to ensure consistent data-to-lead field scale ratios for both. Four scaling parameters (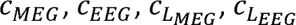) were used to accomplish this, maintaining the relations: 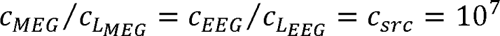, which ensured that estimated source signals for MEG and EEG would have a common scale, and could be transformed into biophysically plausible estimates by dividing by *C_src_*.

Five sets of these scaling parameters were devised to adjust the balance between MEG and EEG data labels, as outlined in Table S1. The first set of scaling parameters was selected based on previous single-modality studies (O’Reilly, Zhu, et al., 2023; Srivastava et al., 2023) to avoid vanishing gradients. However, these produced an imbalance between MEG and EEG data, with a ratio of mean absolute values 〈|***Y**_MEG_*|〉/〈|***Y**_EEG_*|〉 = 1.06/0.541. The second set of scaling parameters was selected to partly redress this imbalance, producing a ratio of 〈|***Y**_MEG_*|〉/〈|***Y**_EEG_*|〉 = 1.06/0.811. The third set was calculated to balance this ratio, i.e., 〈|***Y**_MEG_*|〉/〈|***Y**_EEG_*|〉 = 1. The fourth set was calculated to balance the sum of MEG and EEG absolute values, i.e.,∑|***Y**_MEG_*|∑|***Y**_EEG_*| = 1. The fifth was calculated to balance the standard deviation of MEG and EEG data labels, i.e., σ_MEG_/σ_EEG_ = 1. While these five sets of scaling parameters influenced the performance of RNN models in terms of comparison between ground-truth and reconstructed ERF and ERP waveforms (Figure S4), the relatively small variation did not provide strong evidence for preferring source signals estimated using either one set of scaling values. Weighting EEG more than MEG labels (scaling set 4) biased the model towards the former, enhancing the overall correlation between ground truth and reconstructed waveforms; however, this evaluation is confounded by higher inter-channel correlation for EEG than MEG (Figure S5). Therefore, to calculate the final estimated source signals from the RNN method, those produced by all five sets of scaling values and five random seeds (i.e., 25 estimates) were averaged. ERF and ERP waveforms reconstructed from these RNN source estimates are plotted in Figure 2.

**Figure 2.**
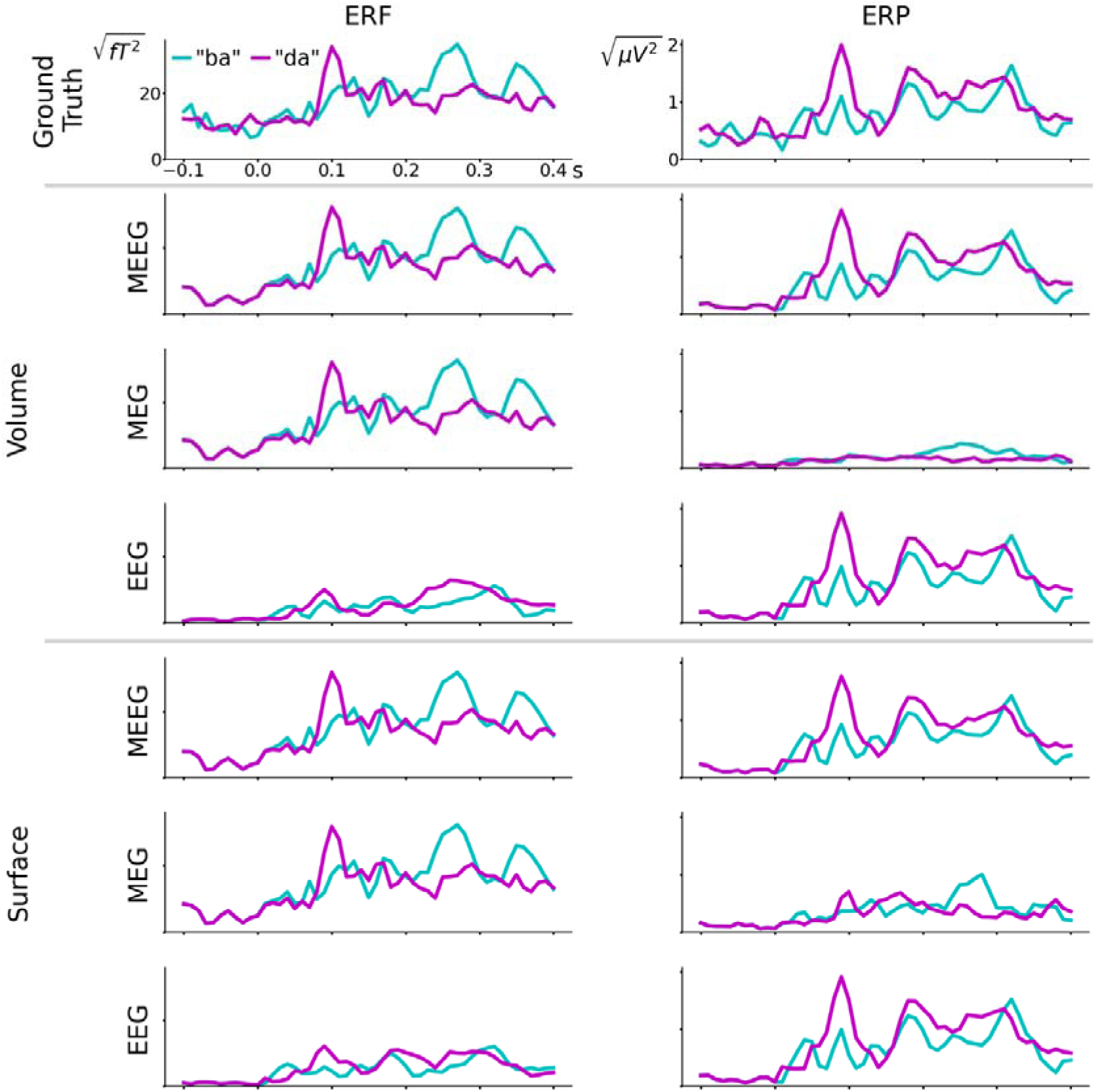
ERF and ERP waveforms reconstructed from source signals estimated from RNN. Root-mean-squared amplitudes across all MEG or EEG channels (for ERF or ERP waveforms, respectively) are plotted on left and right columns. Y-axis scales are the same for each row, and x-axis scales are the same for all plots. Results from volume and surface source spaces are comparable. When fitted to unimodal data (MEG or EEG), reconstructed waveforms for the alternate modality are comparatively poor; consistent with results from eLORETA and MxNE presented in Figure S6 and Figure S7. Analyses of these waveforms are provided in Table 1.

**Table 1.** Evaluation of reconstructed ERF and ERP waveforms and source signals estimated using different methods, source spaces, and fitting data.

| Method | Source space | Fitted to | ERF |  |  | ERP |  |  | SRC<sup>1</sup> |
| --- | --- | --- | --- | --- | --- | --- | --- | --- | --- |
|  |  |  | r | MSE | SNR | r | MSE | SNR | SNR |
| RNN | Volume | MEEG | 0.929 | 4.92e-29 | 10.6 | 0.935 | 1.12e-13 | 15.8 | 13 |
|  |  | MEG | 0.932 | 4.67e-29 | 10.2 | 0.0908 | 8.15e-13 | 11.3 | 10.9 |
|  |  | EEG | -0.00759 | 3.81e-28 | 16.8 | 0.949 | 8.36e-14 | 16.4 | 15.8 |
|  | Surface | MEEG | 0.921 | 5.5e-29 | 10.7 | 0.937 | 1.08e-13 | 15.4 | 13.5 |
|  |  | MEG | 0.927 | 5.06e-29 | 10.2 | 0.301 | 7.52e-13 | 12 | 12.2 |
|  |  | EEG | 0.254 | 3.25e-28 | 17.1 | 0.952 | 7.81e-14 | 16.6 | 16 |
| eLORETA | Volume | MEEG | 0.838 | 1.04e-28 | 6.66 | 0.851 | 2.36e-13 | 6.89 | 5.23 |
|  |  | MEG | 0.862 | 8.9e-29 | 6.48 | 0.0476 | 3.05e-12 | 5.24 | 5.07 |
|  |  | EEG | -0.0997 | 4.37e-28 | 5.2 | 0.889 | 2e-13 | 7.08 | 5.08 |
|  | Surface | MEEG | 0.83 | 1.09e-28 | 6.83 | 0.805 | 3e-13 | 7.41 | 5.82 |
|  |  | MEG | 0.882 | 7.73e-29 | 6.77 | -0.0385 | 3.42e-12 | 3.97 | 4.76 |
|  |  | EEG | 0.262 | 3.28e-28 | 4.82 | 0.836 | 2.92e-13 | 7.6 | 5.47 |
| MxNE | Volume | MEEG | 0.834 | 1.12e-28 | 6.41 | 0.833 | 2.53e-13 | 7.16 | 6.76 |
|  |  | MEG | 0.857 | 1.01e-28 | 6.31 | 0.0177 | 1.38e-12 | 5.52 | 6.83 |
|  |  | EEG | 0.00767 | 7.63e-28 | 5.9 | 0.916 | 1.32e-13 | 6.95 | 5.18 |
|  | Surface | MEEG | 0.757 | 1.48e-28 | 6.71 | 0.754 | 3.54e-13 | 8.54 | 6.86 |
|  |  | MEG | 0.843 | 1.07e-28 | 6.64 | 0.126 | 1.63e-12 | 6.1 | 6.44 |
|  |  | EEG | 0.199 | 3.58e-28 | 3.81 | 0.755 | 3.74e-13 | 8.11 | 6.29 |

<sup>1</sup>SRC: source
Gray shading is applied to highlight performance on the ERF or ERP waveform from the signal modality that was not used when deriving source estimates.

Lead field matrices *L_MEG_* ∈ ∑^160×*n*^_*src*_ and *L_EEG_* ∈ ∑^61×*n*^_*src*_ were used to fit MEG and EEG data, respectively, where *n_src_* = 150 (50 sources × 3 directions) for volume source space, and 132 for surface source space. Lead field matrices *L_MEG_* and *L_EEG_* were concatenated for MEEG; i.e., *L_MEEG_* = [*L_MEG_, L_EEG_*]. Output layer weights (***W**_out_*) in Figure 1 were constructed from scaled lead field matrices, 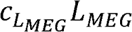 and 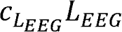, and held fixed during training. Scaled data, *C_MEG_**Y**_MEG_* and *C_EEG_**Y**_EEG_* were provided as labels for supervised learning; i.e., ***Y*** = [*C_MEG_**Y**_MEG_, C_EEG_**Y**_EEG_*] ∈ ∑^197×51×221^ for MEEG data, ***Y*** = *C_MEG_**Y**_MEG_* ∈ ∑^197×51×160^ for MEG data, and ***Y*** = *C_EEG_**Y**_EEG_* ∈ ∑^197×51×61^ for EEG data. In each case, estimated source signals were extracted from the fourth recurrent layer hidden unit activations (i.e., 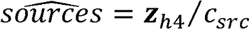.

### 2.5. eLORETA and MxNE

Two established inverse methods were computed using averaged ERF and ERP responses to “ba” and “da” events. Functions in MNE-python were used to implement these. Noise sample covariance was estimated from the pre-stimulus baseline, and data sample covariance was estimated from the post-stimulus period of single-trial data. For the volume source space, the *depth* weight prior was 0.8, and the variance of source components tangential to the cortical surface was weighted by the *loose* parameter 1.0 for free orientation. There was no *depth* prior weighting for the surface source space, and the *loose* parameter was set to zero for fixed orientation.

Exact low-resolution electromagnetic tomography (eLORETA) is claimed to have zero localization error for EEG inverse problems under noiseless conditions (Pascual-Marqui et al., 2011). In reality there is no such ideal situation, but eLORETA is widely used and regarded highly among linear inverse methods for EEG and MEG source reconstruction (Pascual-Marqui et al., 2018). The MNE-python implementation of eLORETA was applied with the `apply_inverse` function. Recommended default parameters were used, and the *lambda2* regularization parameter was set to 1/3^2^, where 3 was the estimated signal-to-noise ratio.

Mixed-norm estimates (MxNE) combine L1 and L2 norm constraints while solving the inverse problem, yielding sparse solutions (Gramfort et al., 2012). We selected MxNE to compare with the RNN approach because both methods incorporate L1 regularization of estimated source signals. It was implemented using the `mixed_norm` function in MNE-python with the regularization parameter *alpha* 5.0; otherwise, recommended default parameters were selected.

### 2.6. Source ranking analysis

Estimated source signals were ranked in reverse order by correlating their projected ERF and ERP waveforms with the ground truth. Individual source signals were multiplied by their lead field coefficients to project them onto sensor space. Then these projections were compared with ground-truth ERF and ERP waveforms using Pearson’s correlation coefficient. For each volume source, estimated signals for three direction components were multiplied by their corresponding lead field coefficients to generate projected waveforms. For both source spaces, each vertex received nine rankings [i.e., from three methods (RNN, eLORETA, MxNE) applied to three data types (MEEG, MEG, EEG)]. These aggregated rankings were used to evaluate the strength of evidence for each vertex’s involvement in generating ERF and ERP waveforms. Only the two most highly ranked sources are plotted (Figure 3 and Figure 4), although all are available from an online repository.

**Figure 3.**
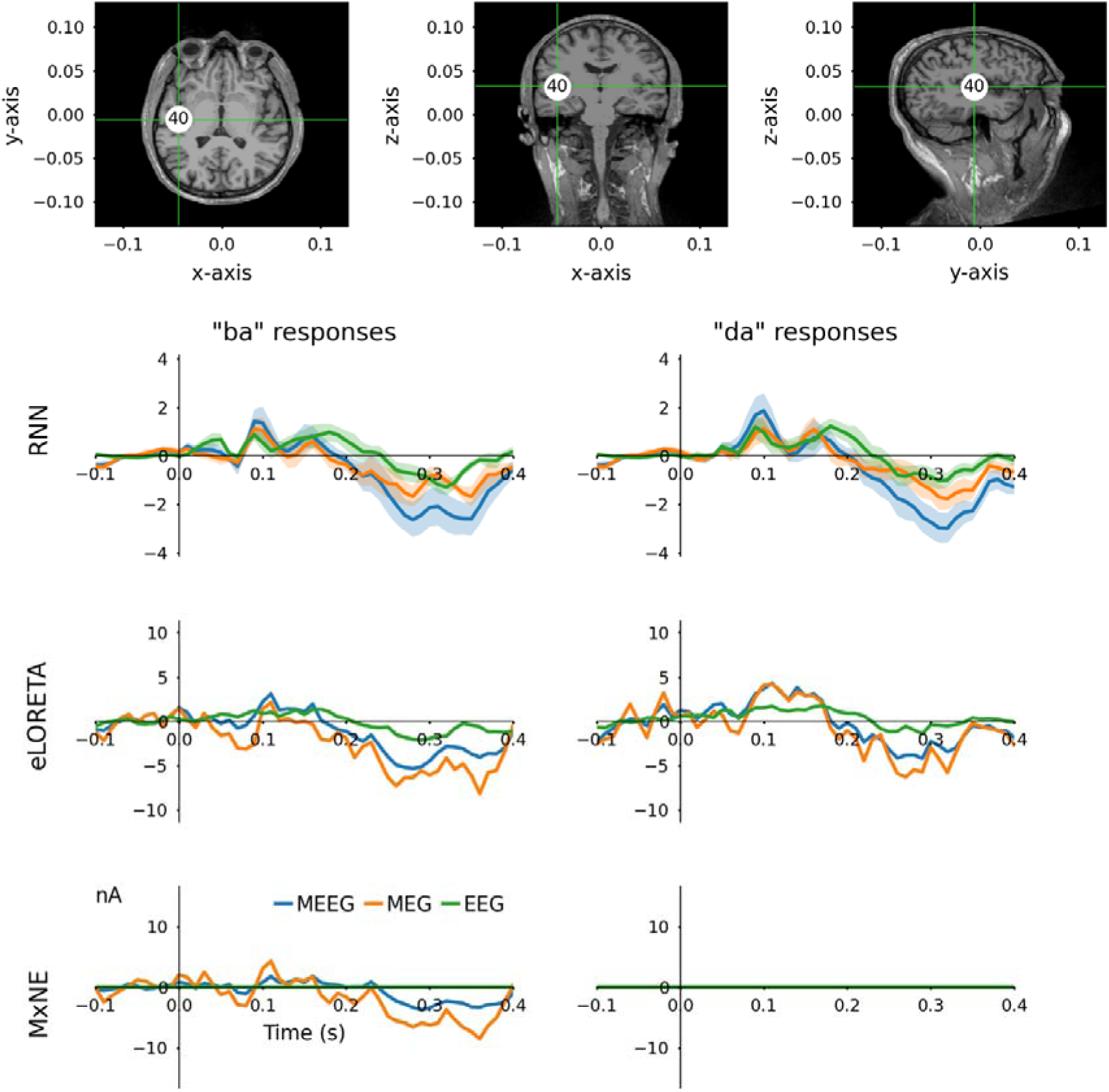
Estimated signals from vertex 40 of the surface source space with fixed dipole orientation normal to the cortical surface. When projected on to MEEG channels using the lead-field matrix, this source was ranked among the top five in terms of correlation between real and projected MEEG for 8 out of 9 source estimates (i.e., three methods [RNN, eLORETA, MxNE] times three data types [MEEG, MEG, EEG]). This source is located in the left temporal lobe, consistent with regions involved in sound and language processing. Shaded error bars represent the standard deviation from 25 source estimates obtained with the RNN method. Different y-axis scales are used for plotting estimated source signals from different methods.

**Figure 4.**
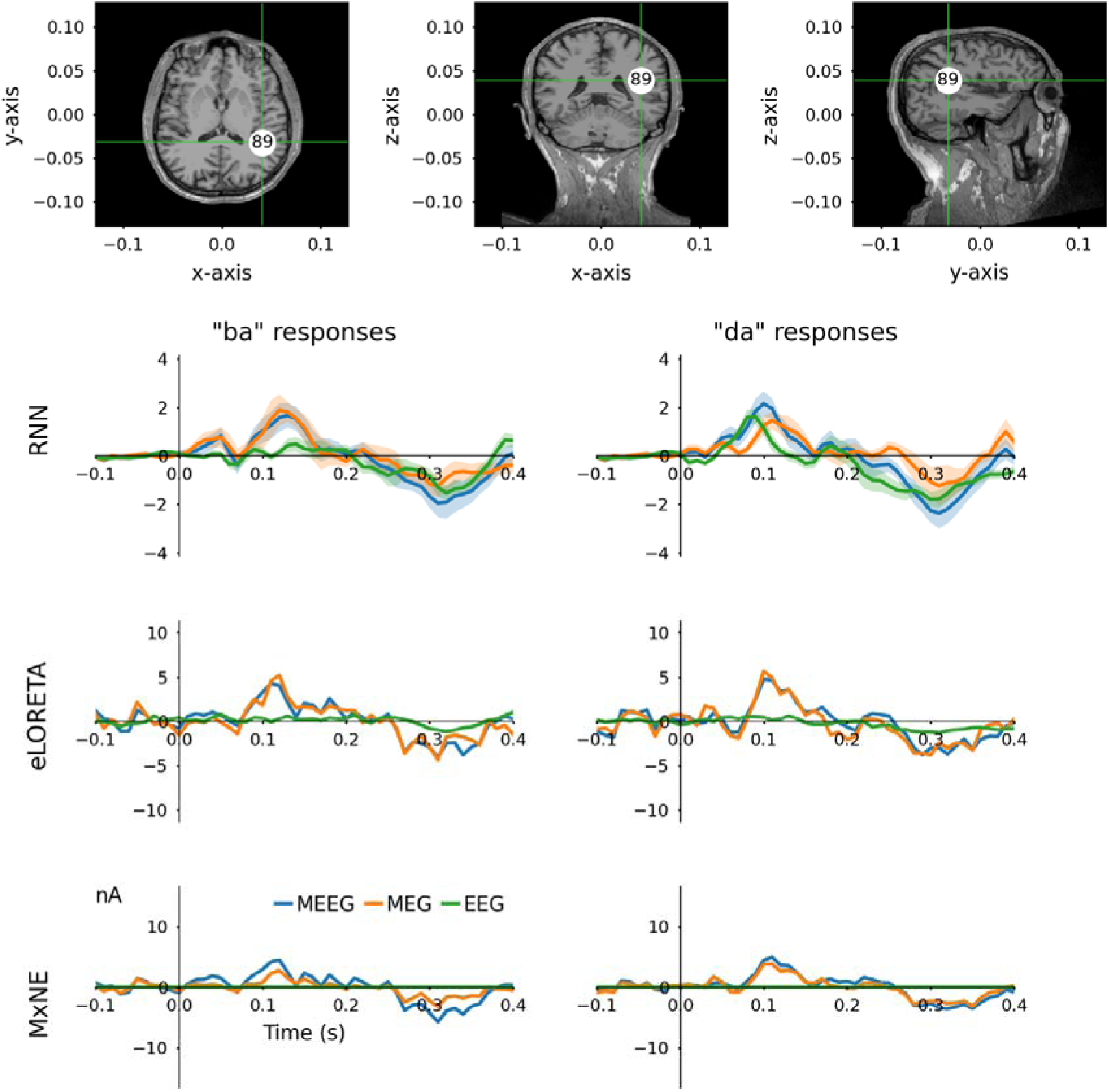
Source signals estimated from vertex 89 of the fixed-orientation surface source space. This source also ranked within the top five with the highest correlation between real and projected MEEG for 8 out of 9 source estimates. Situated in the right posterior temporal lobe, this is consistent with expectations for auditory processing of syllable sounds. Standard deviations of RNN source estimates are denoted with shared error bars. Y-axes scales are different for each method.

The consistency of signals estimated by the RNN method was also studied. Average correlations among each source signal component estimated by 25 RNN models (five sets of scaling parameters, five random seeds) after training without (step 1) and with L1-norm regularization (step 2) were evaluated for each data type (MEEG, MEG, EEG) and source space (volume, surface). For this analysis, three direction components for volume source space vertices were treated separately. Distributions of these average correlations are plotted in Figure 5.

**Figure 5.**
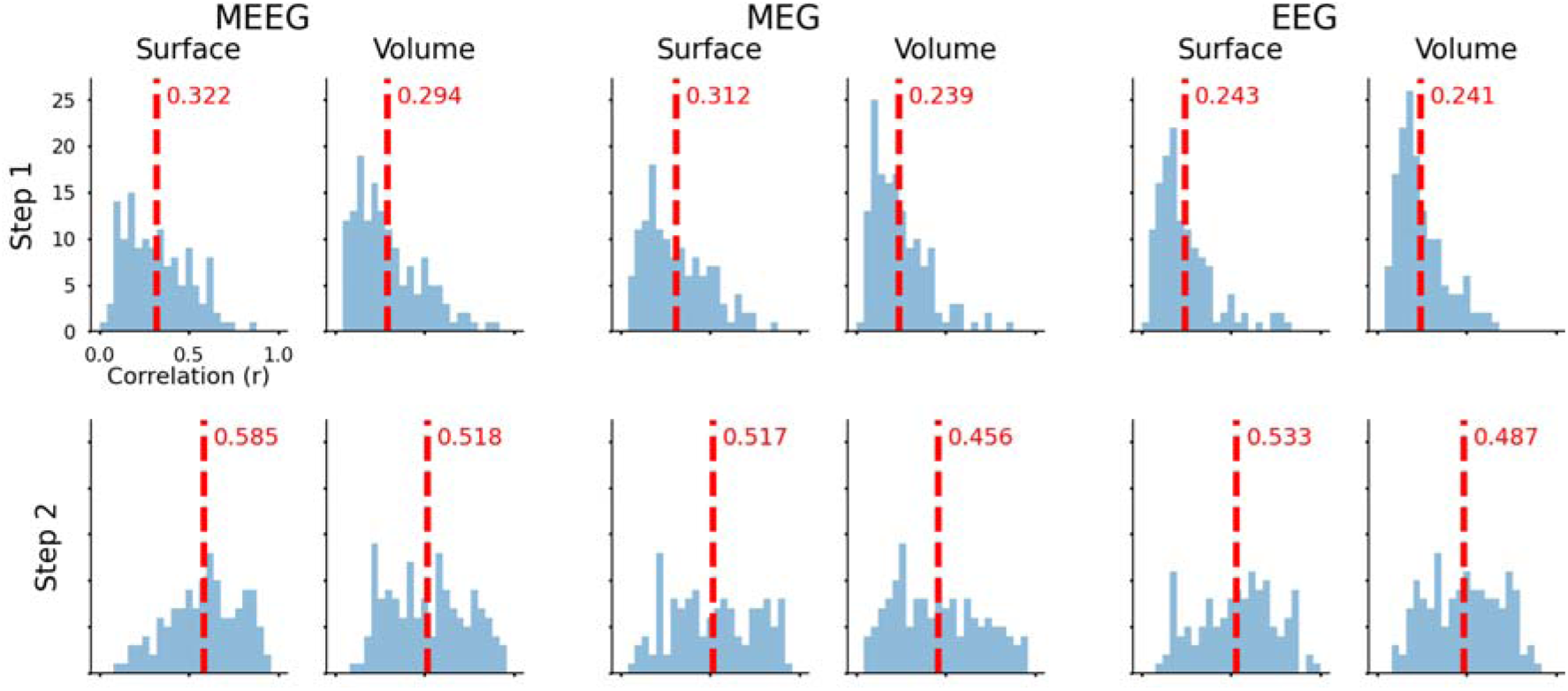
Correlation among source estimates from 25 RNNs (five scales, five seeds) fitted to MEEG, MEG, or EEG data, with surface or volume source spaces. Directions (x, y, z) for volume space sources were treated separately, yielding 150 estimated source signals; the surface source space with fixed orientation had 132 estimated source signals. Step 2 source estimates were more consistent (higher r) than step 1. Surface space solutions are more consistent than those of the volume space, although this could be explained by fewer estimates in the surface space constraining the range of possible solutions. Moreover, MEEG sources were more consistent than single-modality estimates. Means are indicated by vertical dashed lines and reported in red text.

### 2.7. Data analysis and statistical tests

Reconstructed ERF and ERP waveforms were compared with ground-truth waveforms using Pearson’s correlation coefficient (r) and mean squared error (MSE). Signal to noise ratio (SNR) was calculated in decibels (dB) using the formula SNR = 20log_10_(signal_RMS_/noise_RMS_), where signal_RMS_ was the root-mean-squared (RMS) amplitude from the post-stimulus period (>0.0 s) and noise_RMS_ was the RMS amplitude in the pre-stimulus period (<0.0 s). The SNR of reconstructed waveforms and estimated source signals (SRC SNR) were evaluated. These data are reported in Table 1.

Non-parametric statistical hypothesis tests were performed to assess whether differences among samples were statistically significant because they either did not have equal variance and/or did not have a normal distribution. The Brown-Forsythe test was used to check for equal variance (Brown & Forsythe, 1974), and D’Agostino and Pearson’s method was used to test for normality (D’Agostino & Pearson, 1973). Paired samples were tested with Wilcoxon signed-rank tests (Wilcoxon, 1945) and independent samples with Mann-Whitney U-tests (Mann & Whitney, 1947), with two-sided alternatives. Descriptive statistics are reported for variables enclosed in parenthesis, e.g., (sample size *n*, mean *μ*, standard deviation *σ*, median *M*, interquartile range *IQR*), and inferential statistics are reported in square brackets, e.g., [statistic, p-value]. P-values below 0.05 may be considered statistically significant.

### 2.8. Software

Python 3 with matplotlib 3.5.2 (Hunter, 2007), meegkit 0.1.3, mne 1.4.0 (Gramfort et al., 2013), numpy 1.25.2 (Harris et al., 2020), pyvista 0.37.0, scipy 1.11.2 (Virtanen et al., 2020), and tensorflow 2.14.0 (Abadi et al., 2016) were used to conduct this research.

## 3. Results and discussion

### 3.1. Scaling labels influences the correlation between reconstructed and ground-truth ERF and ERP waveforms

Scaling MEG and EEG data labels and their corresponding lead field matrices was necessary to avoid vanishing gradients when training RNNs (Bengio et al., 1994; Pascanu et al., 2013). When label values are very small (***y*** « 1), error gradients propagated backwards through the layers to update network parameters become miniscule (i.e., they “vanish”). Therefore, labels must be scaled appropriately for effective training. This requirement for modelling event-related neural signals with RNNs has been highlighted previously (O’Reilly, Wehrman, et al., 2022). In the present study, additional consideration was given to balancing the amplitudes of labels derived from MEG and EEG because they have different raw signal magnitudes.

Different scaling coefficients for MEG and EEG data labels and lead-field matrices influenced the performance of RNNs, as detailed in Table S1. The average source estimates from five RNNs (i.e., 5 random seeds) trained using each set of scaling values were evaluated by correlation between reconstructed and ground-truth ERF and ERP waveforms. These correlations are plotted in Figure S4. The scaled MEG to EEG standard deviation (σ_MEG_/σ_EEG_) ratio positively correlated with the ERF to ERP correlation (r_ERF_/r_ERP_) ratio from RNNs trained with MEEG. In contrast, the overall correlation of combined ERF and ERP waveforms (r_both_) from RNNs trained with MEEG had an inverse relationship with σ_MEG_/σ_EEG_, indicating that the correlation with the ERP has more influence over r_both_. Models trained with single modality labels are consistent with these observations. Outputs from RNNs trained with scaled MEG labels only, have a positive relationship between r_ERF_ and σ_MEG_/σ_EEG_. In contrast, outputs from RNNs trained with scaled EEG labels only, have a negative relationship between r_ERP_ and σ_MEG_/σ_EEG_. For RNNs trained with MEG-only or EEG-only, performance appeared to level off towards the favoured side of this ratio: higher for MEG, lower for EEG. The same pattern of results was observed for volume and surface source spaces.

These observations suggest that the overall correlation metric r_both_ is biased towards EEG data because EEG had fewer channels and higher inter-channel correlation (Figure S5) than MEG, making it “easier” for the model to fit and thereby achieve higher overall correlation. Biophysical processes cause EEG channels to have higher inter-channel correlation than MEG channels (Luck, 2014; Nunez & Srinivasan, 2009), so a model trained with larger amplitude labels derived from EEG than MEG appears to perform better at reproducing ERF and ERP waveforms overall. Despite the influence of scaling to sway the performance of RNNs to favour of fitting ERF or ERP waveforms, all of the scaling values used produced outputs that were highly correlated with ground-truth ERF and ERP waveforms. Therefore, source estimates from all 25 RNN models (five scales × five seeds) were averaged to obtain the final source estimates used to evaluate reconstructed ERF and ERP waveforms against ground-truth waveforms for the RNN method reported in section 3.2.

### 3.2. Fewer training epochs are required when fine-tuning models with L1 regularization

Distributions of completed epochs for RNN training steps 1 and 2 are plotted in Figure S3. In every case, training stopped before 3000 epochs were completed. There were no statistically significant differences between volume (n = 150, μ = 1119, σ = 450, M = 1112, IQR = 603) or surface (n = 150, μ = 1167, σ = 425, M = 1150, IQR = 638) source spaces, nor were there among models trained with MEEG (n = 100, μ = 1184, σ = 436, M = 1160, IQR = 652), MEG-only (n = 100, μ = 1156, σ = 422, M = 1134, IQR = 571), or EEG-only (n = 100, μ = 1089, σ = 452, M = 1076, IQR = 584). In training step 2, when fine-tuning the RNN with L1 regularization applied to the source estimation layer (SimpleRNN 4 in Figure 1), fewer epochs were completed (n = 150, μ = 923, σ = 375, M = 884, IQR = 494) than in training step 1 (n = 150, μ = 1363, σ = 384, M = 1346, IQR = 504) [W = 1515.5, p = 7.21e-15]. Observing a statistically significant difference between the number of completed training epochs in step 1 and step 2 is unsurprising, given that deep neural network models generally require less training to converge when pre-trained before transfer learning (Hosna et al., 2022). Adding the L1-norm constraint to source estimation layer activations also produced more consistent source estimates from RNNs (Figure 5), and caused retrograde attenuation of hidden unit activations in preceding layers (e.g., as shown in Figure S8).

### 3.3. Reconstructed ERF and ERP waveforms from RNN are “better” than eLORETA and MxNE

Ground-truth ERF and ERP waveforms are plotted on the top row of Figure 2 above waveforms reconstructed from the RNN method. Source estimates from RNNs fitted to MEEG data reconstructed ERF and ERP waveforms with high fidelity. In contrast, source signals estimated from RNNs trained with MEG-only or EEG-only produced comparatively poor reconstructions of contra waveforms; i.e., those from RNNs trained with EEG-only had poor ERF reconstructions, whereas those from RNNs trained with MEG-only had poor ERP reconstructions. Comparable plots for eLORETA and MxNE are in Figure S6 and Figure S7, respectively, where a similar pattern is observed. Analyses of reconstructed ERF and ERP waveforms are reported in Table 1. An example of signals generated by each hidden unit of the RNN is plotted in Figure S8. These signals generated internally by the model are analogous to neurophysiological signals generated by the brain that produce ERF and ERP waveforms. This functional distinction between the RNN method and conventional linear inverse solutions, such as eLORETA and MxNE, offers the possibility of using the RNN as a “model organism” for investigating computational mechanisms associated with event-related neural signal generation (Lindsay & Bau, 2023; O’Reilly, 2022a; O’Reilly, Angsuwatanakul, et al., 2022).

Metrics calculated for reconstructed ERF and ERP waveforms are provided in Table 1. There were no statistically significant differences in r, MSE, or SNR between volume (n = 18; r: μ = 0.594, σ = 0.416, M = 0.844, IQR = 0.851; MSE: μ = 3.48e-13, σ = 7.42e-13, M = 4.18e-14, IQR = 2.27e-13; SNR: μ = 8.72, σ = 3.81, M = 6.92, IQR = 4.2) and surface source spaces (n = 18; r: μ = 0.628, σ = 0.326, M = 0.781, IQR = 0.6; MSE: μ = 4.06e-13, σ = 8.3e-13, M = 3.91e-14, IQR = 3.41e-13; SNR: μ = 8.85, σ = 3.96, M = 7.51, IQR = 3.92) [r: W = 82.0, p = 0.899, MSE: W = 62.0, p = 0.325, SNR: W = 53.0, p = 0.167]. Metrics for waveforms reconstructed from source signals estimated by the RNN (n = 12; r: μ = 0.677, σ = 0.373, M = 0.928, IQR = 0.646; MSE: μ = 1.62e-13, σ = 2.81e-13, M = 3.91e-14, IQR = 1.09e-13; SNR: μ = 13.6, σ = 2.82, M = 13.7, IQR = 5.75) were statistically significantly better, meaning higher r and SNR and lower MSE, from those of eLORETA (n = 12; r: μ = 0.58, σ = 0.389, M = 0.833, IQR = 0.646; MSE: μ = 6.25e-13, σ = 1.18e-12, M = 9.98e-14, IQR = 2.94e-13; SNR: μ = 6.25, σ = 1.1, M = 6.71, IQR = 1.71) [r: W = 1.0, p = 0.000977, MSE: W = 0.0, p = 0.000488, SNR: W = 0.0, p = 0.000488] and MxNE (n = 12; r: μ = 0.575, σ = 0.351, M = 0.756, IQR = 0.656; MSE: μ = 3.44e-13, σ = 5.41e-13, M = 6.58e-14, IQR = 3.59e-13; SNR: μ = 6.51, σ = 1.16, M = 6.52, IQR = 0.956) [r: W = 1.0, p = 0.000977, MSE: W = 0.0, p = 0.000488, SNR: W = 0.0, p = 0.000488], whereas those from eLORETA and MxNE were not statistically significantly different [r: W = 27.0, p = 0.380, MSE: W = 32.0, p = 0.622, SNR: W = 25.0, p = 0.301].

The signal to noise ratio of estimated source signals (SRC SNR) was not statistically significantly different between volume (n = 9, μ = 8.21, σ = 3.81, M = 6.76, IQR = 5.76) and surface (n = 9, μ = 8.6, σ = 3.92, M = 6.44, IQR = 6.39) source spaces [W = 7.0, p = 0.0742]. However, they were statistically significantly different between RNN (n = 6, μ = 13.6, σ = 1.84, M = 13.3, IQR = 2.85) and eLORETA (n = 6, μ = 5.24, σ = 0.337, M = 5.16, IQR = 0.342) [W = 0.0, p = 0.03125], RNN and MxNE (n = 6, μ = 6.39, σ = 0.582, M = 6.6, IQR = 0.482) [W = 0.0, p = 0.03125], and eLORETA and MxNE [W = 0.0, p = 0.03125]. RNN and MxNE methods incorporate an L1 norm constraint, producing higher SRC SNR than eLORETA. However, MxNE does this by turning off sources, whereas the RNN does not eliminate any sources. Training RNNs with input signals that go high only during event-on time points (bottom of Figure 1) induces common baseline activity for both event types because there is no distinction between input representations before stimulus onset. Therefore, baseline activity estimated by the RNN converges towards the mean of both event types and is less than that estimated by eLORETA or MxNE methods, producing higher SNR for estimated source signals and reconstructed waveforms. This can be viewed as an advantage of the RNN method with the assumption that there are no systematic differences in preparatory activity or overlap from previous events between event types that would cause their baseline activity to be different.

These results demonstrate that the RNN method can produce source signals that effectively reconstruct event-related neural signals observed from simultaneously recorded MEG and EEG. These reconstructed ERF and ERP waveforms had higher correlation and lower MSE evaluated against ground-truth ERF and ERP waveforms and higher SNR compared with those derived from source estimates produced by eLORETA or MxNE methods (Table 1). All methods applied to MEEG data produced source signals better at reconstructing both ERF and ERP waveforms compared with single modality data alone. As seen from Figure 2 (and Figure S6 and Figure S7), source estimates derived from MEG-only or EEG-only were comparatively poor at reconstructing event-related signals from the modality not used to derive those source estimates. This highlights that MEG and EEG sensitivity to different sources varies according to the location and orientation of those sources relative to the electrode or magnetometer (Nunez & Srinivasan, 2009). Electric potential gets smeared through the skull and the magnetic field does not. Moreover, MEG and EEG are sensitive to perpendicular components of the electromagnetic field and have different sensor geometry relative to the brain. As such, it is not unexpected that source signals estimated using either modality alone are considerably different (e.g., see Figure 3 and Figure 4).

### 3.4. Source rankings from surface source space provide stronger evidence for vertex relevance

From three source estimation techniques (RNN, eLORETA, and MxNE) applied to three data types (MEEG, MEG, and EEG), the top five highest-ranked sources based on the correlation between their projections and ground-truth ERF and ERP waveforms are reported in Table 2. From volume source space, vertices 21 and 23 were most consistently ranked in the top five across estimation methods and data types, at 6/9 and 5/9, respectively. The probability of any single volume-space vertex being ranked in the top five by random chance would be 5/50 = 0.1, and if each observation of top-five rankings was independent, the recurring probability would be 0.1^n^ for *n* repetitions; therefore, the chance of vertices 21 and 23 appearing in 6/9 and 5/9 of the top-five ranked source estimates would be 10^−6^ and 10^−5^. From the surface source space, vertices 40 and 89 were ranked among the top five of 8/9 source estimates. By random chance, the probability of an individual vertex in the surface source space being ranked in the top five would be 5/132 = 0.0378; therefore, the probability of observing these results for vertices 40 and 89 is 0.0378^8^ = 4.168×10^−12^.

**Table 2.** Source signals ranked by correlation between projected and observed ERF and ERP waveforms.

| Source space<sup>1</sup> | Method | Fitted to | Ranked source estimates<sup>2</sup> (vertex number: $r_{\text{both}}$)<sup>3</sup> | | | | |
| --- | --- | --- | --- | --- | --- | --- | --- |
|  |  |  | #1 | #2 | #3 | #4 | #5 |
| Volume | RNN | MEEG | 22: 0.737 | 35: 0.706 | 23: 0.703 | 36: 0.66 | 17: 0.657 |
|  |  | MEG | 16: 0.301 | 23: 0.269 | 24: 0.266 | 17: 0.254 | 37: 0.222 |
|  |  | EEG | 32: 0.805 | 35: 0.801 | 21: 0.784 | 22: 0.774 | 36: 0.772 |
|  | eLORETA | MEEG | 22: 0.689 | 35: 0.543 | 21: 0.45 | 18: 0.425 | 23: 0.389 |
|  |  | MEG | 34: 0.294 | 7: 0.289 | 17: 0.286 | 21: 0.234 | 38: 0.205 |
|  |  | EEG | 22: 0.736 | 35: 0.677 | 18: 0.605 | 11: 0.598 | 23: 0.587 |
|  | MxNE | MEEG | 21: 0.472 | 23: 0.466 | 37: 0.315 | 24: 0.304 | 36: 0.279 |
|  |  | MEG | 21: 0.362 | 7: 0.326 | 17: 0.28 | 34: 0.265 | 24: 0.214 |
|  |  | EEG | 10: 0.559 | 21: 0.442 | 27: 0.431 | 48: 0.306 | 12: 0.276 |
| Surface | RNN | MEEG | 89: 0.526 | 63: 0.514 | 115: 0.453 | 2: 0.446 | 40: 0.444 |
|  |  | MEG | 40: 0.384 | 89: 0.345 | 37: 0.323 | 1: 0.293 | 116: 0.284 |
|  |  | EEG | 40: 0.623 | 28: 0.588 | 89: 0.583 | 76: 0.559 | 37: 0.531 |
|  | eLORETA | MEEG | 89: 0.419 | 40: 0.358 | 108: 0.339 | 115: 0.322 | 91: 0.278 |
|  |  | MEG | 89: 0.37 | 40: 0.259 | 116: 0.207 | 22: 0.195 | 35: 0.183 |
|  |  | EEG | 40: 0.485 | 89: 0.482 | 31: 0.403 | 115: 0.383 | 108: 0.382 |
|  | MxNE | MEEG | 89: 0.427 | 115: 0.375 | 116: 0.324 | 2: 0.303 | 40: 0.262 |
|  |  | MEG | 89: 0.405 | 116: 0.254 | 35: 0.218 | 37: 0.206 | 40: 0.187 |
|  |  | EEG | 115: 0.431 | 63: 0.308 | 100: 0.296 | 36: 0.282 | 2: 0.28 |

<sup>1</sup>Volume space projections were computed from three signal components (x, y, and z directions) for each of 50 vertices, whereas surface space projections were computed from individual signal components for 132 vertices.
<sup>2</sup>Ranked from highest to lowest correlation between projected and observed MEEG.
<sup>3</sup> $r_{\text{both}}$ was calculated by comparing projections with combined ERF and ERP waveforms.

The lower probability of observed rankings for vertices 40 and 89 of the surface source space can be viewed as evidence that these estimates are more reliable than those of the volume source space. As such, estimated source signals from surface space vertices 40 and 89 are plotted in Figure 3 and Figure 4, respectively. Vertices 40 and 89 are located in the left and right temporal lobes, respectively, where one might expect syllable sounds to excite neural tissue specialised for sound and language processing (Rong et al., 2018; Schirmer et al., 2012). Source estimates from the RNN are plotted with shaded error bars, quantifying variability in the data, which is impossible for the other two deterministic methods. The amplitude range for RNN source signals is smaller than that of eLORETA and MxNE, and RNN source signals have less pre-stimulus baseline activity. The MxNE method turns off vertices 40 and 89 for both “ba” and “da” responses when fitted to EEG and vertex 40 for “da” responses when fitted to MEEG and MEG as well. Signals from the RNN are qualitatively smoother than those produced by the other methods, reflecting dependence on the sequential structure of internal dynamics (Figure S8), in contrast with eLORETA and MxNE that can be applied to individual time samples. Aside from these differences, the estimated source signals from all three methods at these two vertices are qualitatively similar, indicating that they resemble the likely distribution of the current source density at these locations.

### 3.5. Analysis of individual source projections

Distributions of r, MSE, and SNR measured from waveforms projected from individual estimated sources are plotted in Figure S9, Figure S10, and Figure S11, respectively. Three signal components were projected simultaneously for the volume source space. In contrast, one signal component was projected for each source vertex for the surface source space, reflecting free and fixed dipole orientations, respectively. For r and MSE, projections were compared with combined ground-truth ERF and ERP waveforms, and for calculating SNR, the projected ERF and ERP waveforms were also used.

Correlation between projections and ground-truth ERF and ERP waveforms was higher for volume sources (n = 419, μ = 0.187, σ = 0.257, M = 0.141, IQR = 0.342) than surface sources (n = 1028, μ = 0.101, σ = 0.146, M = 0.0705, IQR = 0.192) [U = 255965.0, p = 1.788e-08]; probably because volume sources had three projected signal components, whereas surface sources had only one projected signal component. Source estimates from the RNN (n = 546, μ = 0.211, σ = 0.222, M = 0.196, IQR = 0.295) produced projections that had a higher correlation with ground-truth waveforms than those from eLORETA (n = 546, μ = 0.0758, σ = 0.149, M = 0.0515, IQR = 0.164) [W = 3030.0, p = 4.857e-84] or MxNE (n = 355, μ = 0.0736, σ = 0.133, M = 0.0525, IQR = 0.158) [U = 133944.0, p = 2.988e-22]; however, those of eLORETA and MxNE were not statistically significantly different [U = 93815.0, p = 0.417]. Projections of source estimates derived from EEG (n = 463, μ = 0.229, σ = 0.198, M = 0.202, IQR = 0.271) had higher correlation than those derived from MEEG (n = 503, μ = 0.139, σ = 0.173, M = 0.104, IQR = 0.202) [U = 84644.0, p = 2.127e-13] or MEG (n = 481, μ = 0.0131, σ = 0.123, M = 0.00555, IQR = 0.151) [U = 37265.0, p = 4.936e-70]. The same pattern of statistical significance was observed for MSE, but with inverted relationships; e.g., volume space projections had higher r and lower MSE than surface space projections.

The finding that source estimates obtained by fitting EEG-only produced projections with higher overall correlation can be explained by higher inter-channel correlation of EEG (n = 876, μ = 0.538, σ = 0.275, M = 0.553, IQR = 0.459) compared with MEG (n = 6295, μ = 0.346, σ = 0.248, M = 0.291, IQR = 0.374) [U = 1665455.0, p = 1.230e-80]. Distributions of non-zero inter-channel correlation coefficients from MEG and EEG are plotted in Figure S5. There were 61 EEG channels, making 61^2^ – 61 = 1830 unique pairwise comparisons between channels, of which 876/1830 = 47.9 % were non-zero. For MEG, 160 channels gave 160^2^ – 160 = 12720 unique pairwise comparisons, where 6295/12720 = 49.5 % were non-zero. EEG’s higher average inter-channel correlation reflects a greater field spread than MEG as it passes through the skull (Luck, 2014; Nunez & Srinivasan, 2009). This tendency for EEG channels to have higher correlation, and EEG amplitudes being orders of magnitude greater than MEG, causes r and MSE metrics applied to combined ERF and ERP waveforms to be biased towards the latter.

In terms of projection SNRs, volume sources (n = 419, μ = 7.92, σ = 4.66, M = 6.5, IQR = 6.74) were lower than surface sources (n = 1028, μ = 8.74, σ = 5.52, M = 7.46, IQR = 7.77) [U = 199267.0, p = 0.0255]. Projections from source signals estimated using the RNN method (n = 546, μ = 14, σ = 3.97, M = 13.9, IQR = 5.21) had higher SNR than those of eLORETA (n = 546, μ = 5.01, σ = 2.45, M = 5.01, IQR = 2.99) [W = 96.0, p = 6.617e-91] or MxNE (n = 355, μ = 5.49, σ = 2.67, M = 5.49, IQR = 3.57) [U = 187110.0, p = 1.912e-123], whereas those from MxNE had higher SNR than those of eLORETA [U = 87648.0, p = 0.0152]. The SNRs of projections from source estimates derived from MEEG (n = 503, μ = 8.55, σ = 5.28, M = 7, IQR = 7.21) were not statistically significantly different from those of MEG (n = 481, μ = 7.67, σ = 4.49, M = 6.79, IQR = 6.05) [U = 129501.0, p = 0.0556] or EEG (n = 463, μ = 9.32, σ = 5.93, M = 7.82, IQR = 9.68) [U = 108850.0, p = 0.0796]. However, those from EEG were statistically significantly higher than MEG [U = 96162.0, p = 0.000287].

### 3.6. Training RNNs with MEEG labels and L1 regularization produced more consistent source estimates

Distributions of mean source signal correlation among estimates from 25 RNN models are plotted in Figure 5. The upper triangle of each cross-correlation matrix of 25 estimates (n = 300) was averaged to find the mean correlation for each source signal component in volume and surface source spaces. Step 2 (n = 846, μ = 0.514, σ = 0.223, M = 0.519, IQR = 0.358) had higher correlations than step 1 (n = 846, μ = 0.274, σ = 0.171, M = 0.228, IQR = 0.221) models [W = 6928.0, p = 1.296e-129], demonstrating that the L1 constraint for fine-tuning produces more consistent source signal estimates. The surface source space (n = 792, μ = 0.419, σ = 0.236, M = 0.384, IQR = 0.391) had higher mean correlations than the volume source space (n = 900, μ = 0.372, σ = 0.226, M = 0.323, IQR = 0.34) [U = 397015.0, p = 5.121e-05]; potentially because the domain of all possible solutions constrained by the surface source space with fixed orientation is smaller than the domain of solutions constrained by the volume source space (Henson, Mattout, et al., 2009).

Source estimates from RNNs trained with MEEG (n = 564, μ = 0.428, σ = 0.233, M = 0.409, IQR = 0.38) had higher mean correlations than those from RNNs trained with MEG-only (n = 564, μ = 0.379, σ = 0.234, M = 0.332, IQR = 0.351) [W = 50824.0, p = 9.396e-14] or EEG-only (n = 564, μ = 0.375, σ = 0.226, M = 0.326, IQR = 0.367) [W = 57817.0, p = 1.672e-08]. However, the difference between mean source signal correlations across 25 RNNs trained with MEG-only or EEG-only was not statistically significantly different [W = 76958.0, p = 0.484]. These observations could be interpreted in light of the additional constraints placed on the solution space by combined MEEG, which produces more consistent source estimates than either modality individually by narrowing the search space of possible solutions (Chowdhury et al., 2015, 2018; Dale et al., 1993). This type of analysis cannot be applied to eLORETA or MxNE because they are deterministic.

### 3.7. Limitations

Forward conduction models were computed from a template head BEM model that was morphed using accurate head digitization points. An anatomically accurate head model created from an MRI scan of the subject from whom MEG and EEG data were collected would have been preferable (Henson, Mattout, et al., 2009). Using a template head model does not inherently invalidate the results from inverse source estimations but warrants an air of caution regarding their over-interpretation (Akalin Acar & Makeig, 2013; Das et al., 2020; Liu et al., 2023). As with all inverse methods applied to real MEG and EEG data, there are no ground-truth measurements to validate current source density estimates; nevertheless, results from the different methods can be aggregated to form a picture of the likely underlying distribution of neural activity that cannot be measured directly.

This study used fewer source space vertices (i.e., 50 for volume and 132 for surface source spaces) than sometimes used with distributed source reconstruction methods, which can reach thousands (Ahlfors & Hämäläinen, 2012). This study’s relatively low number of vertices allowed the RNN to reliably converge to solutions that reproduced the observed ERF and ERP waveforms. Using thousands of source space vertices with the RNN method would require considerably more model hidden units, consuming significantly more computational resources during training. However, this limitation is not necessarily overly restrictive, given that regions of synchronously active cortex that produce MEG and EEG signals measured from outside the head are considered to be approximately equal to or greater than 6 cm^2^ (Nunez & Srinivasan, 2009). The surface source space with 132 vertices had an average distance between vertices of 1.53 cm; therefore, it should manage to locate the sources of ERF and ERP components to a reasonable approximation.

The computational requirements of training RNNs were orders of magnitude greater than that of applying eLORETA or MxNE methods. It took approximately nine hours of computer processing time to train 300 RNN models for this study; i.e., 2 source spaces (volume, surface) × 3 data label types (MEEG, MEG, EEG) × 5 scaling values × 5 random seeds × 2 training steps. In contrast, eLORETA and MxNE took less than a few minutes. Whilst more resources are required to implement the RNN method than other inverse solutions, most modern computing workstations can train an RNN in reasonable timescales for post-hoc analysis of neural signals. Moreover, the advantages of using an RNN for source estimation may outweigh its higher computational costs; specifically, it provides a better reproduction of ERF and ERP waveforms, higher SNR, and a model for investigating event-related signal generation processes.

At most, the RNN is an abstract model of neural signal processing, metaphorically representing biological neural networks that generate ERF and ERP waveforms. It lacks sufficient biological realism to make inference about the underlying neurophysiology. Details of the training algorithm, computational units, network connectivity, and objective of the RNN do not reflect biological reality. Brains develop (“are trained”) through evolution and experience, consisting of genetically and neurochemically heterogeneous processing units, comprise comparatively massive interconnected neural circuits with imprecisely characterised network topologies, and implement the objective of controlling perception and behaviour. In contrast, the RNN has a simple six-layer structure with relatively homogenous computational units (the only difference being rectified vs. linear activation function), optimized by gradient descent to reproduce labels derived from MEG and EEG signals.

## 4. Conclusions

The RNN method is effective at estimating event-related neural source signals from MEEG, outperforming the linear inverse solutions in every way except for computational complexity. It transforms simple time sequences representing physical/cognitive events into complex internal dynamics that effectively replicate ERF and ERP waveforms. Its dependence on sequential relationships among internal signals paints a sharp distinction with linear inverse MEEG source estimation methods applied to individual time points. This reliance on sequential dynamics might explain why estimated source signals from the RNN are smoother and have less baseline noise than those from the other methods.

While the RNN hidden unit activations are not precisely matched to biological neural signals, they demonstrate how a relatively simple representation of psychophysiological events can undergo a hierarchy of transformations to elicit ERF and ERP waveforms. This analogy to human brain function is not available from conventional inverse solutions. It opens up the possibility of using the RNN to study computations underpinning event-related neural signal generation. In future studies, biophysically-inspired constraints may be incorporated to explore how structural and functional changes to the recurrent network influence its performance at source estimation. These constraints may include more diverse patterns of connectivity among units and layers of the RNN (Perich & Rajan, 2020), controlling the relative balance of excitation and inhibition in the network (Hertäg & Clopath, 2022), and employing more sophisticated artificial neuron models.

This study also showed empirically that surface source space with fixed surface-normal orientation is preferable to volume source space with free orientation because surface source space produced more consistent estimates and source rankings. This finding is consistent with biophysical arguments of the likely sources of most ERF and ERP signal components being postsynaptic potentials at the distal or apical dendrites of cortical pyramidal neurons oriented radially within the cortical sheet (Luck, 2014; Nunez & Srinivasan, 2009); with some notable exceptions such as auditory brainstem potentials (Moore, 1987). Moreover, the results indicate that MEEG is superior to MEG or EEG, yielding more consistent source signal estimates by constraining the space of solutions that can fit the observed data.

## Acknowledgements

The authors acknowledge the facilities and technical assistance at the KIT-Macquarie Brain Research (MEG) Laboratory, part of the Australian National Imaging Facility. This work was financially supported by King Mongkut’s Institute of Technology Ladkrabang.

## Data availability

Data and code from this study will be shared publicly via an Open Science Foundation repository.

## Ethical review

The Human Research Ethics Committee at Macquarie University reviewed and approved the experimental protocol.

## Author contributions (CRediT statement)

JAO: Conceptualization, Methodology, Software, Formal Analysis, Data Curation, Writing - original draft, Writing - reviewing and editing, Visualization, Funding acquisition. JDZ: Conceptualization, Methodology, Software, Data Curation, Writing - reviewing and editing. PFS: Conceptualization, Resources, Writing - reviewing and editing, Supervision, Funding acquisition.

## Supplementary material

**Figure S1.**
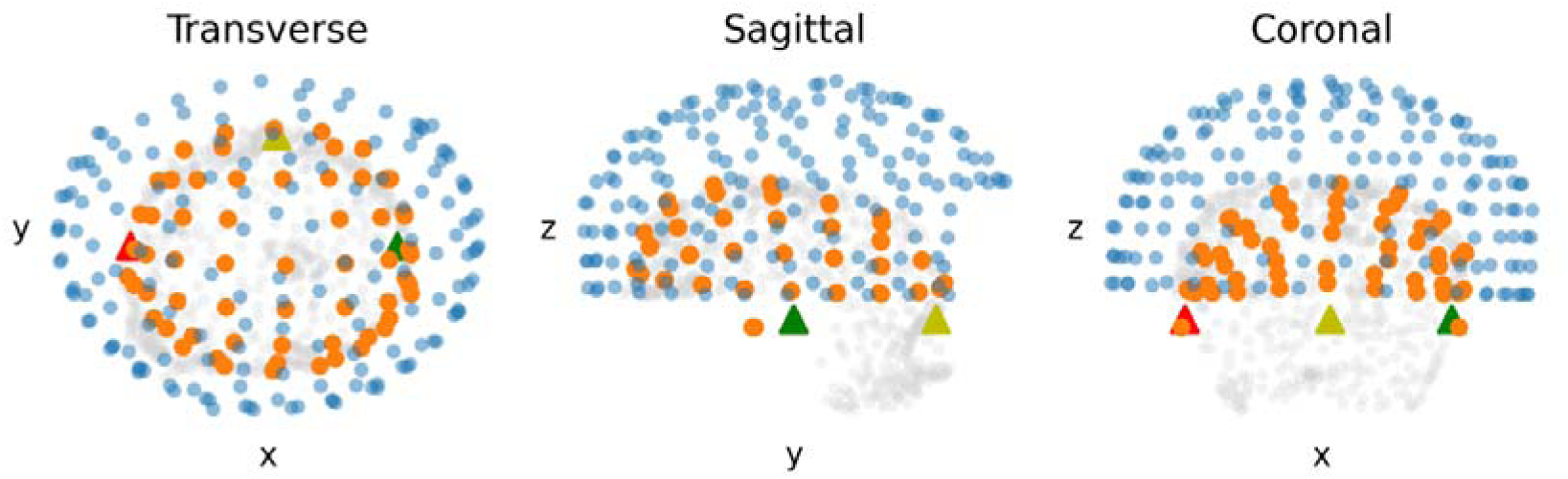
Alignment of EEG and MEG channel locations with head digitization. Locations of left and right preauricular areas and the nasion are marked with red, green, and yellow triangles, respectively; MEG sensor positions are blue circles, EEG sensor positions are orange circles, and head digitization points are light grey circles.

**Figure S2.**
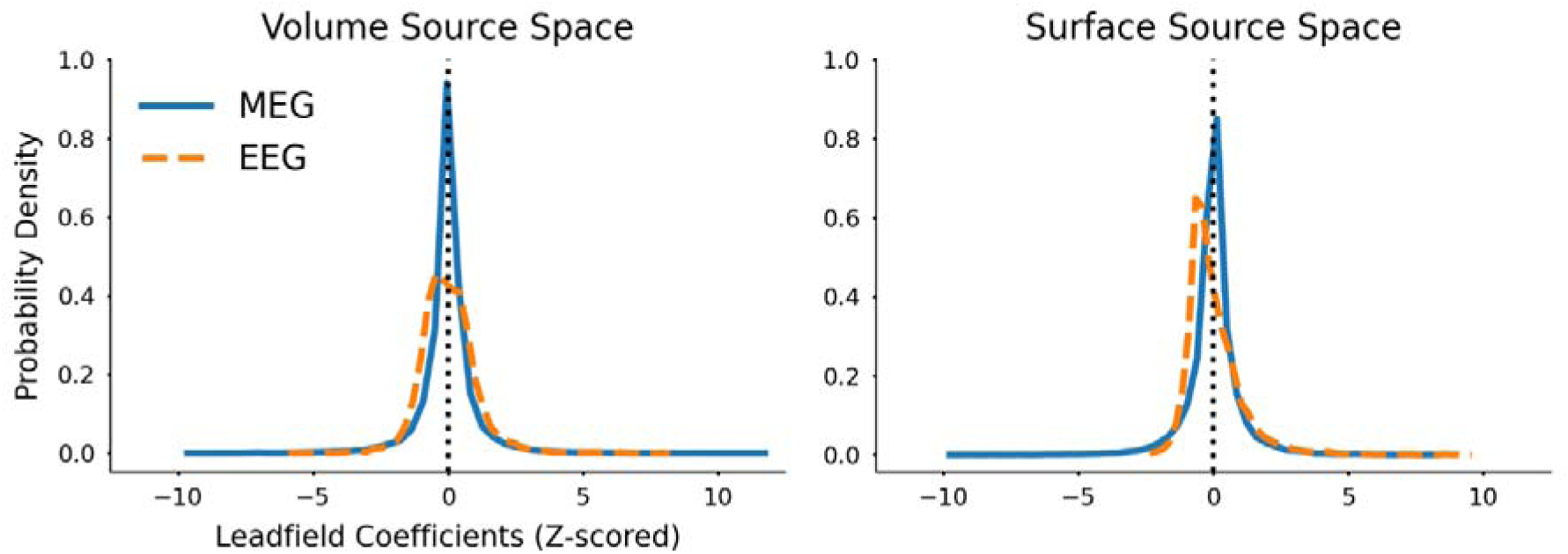
Comparison between distributions of MEG and EEG lead field coefficients from volume and surface source spaces. Each lead-field matrix was standardized to zero mean and unit variance (Z-scored) for visualization. The MEG lead field matrix distributions have higher kurtosis than those from EEG lead field matrices, due to different attenuation of electric and magnetic fields travelling through tissues. The surface source space with fixed orientation normal to the cortical surface caused greater skewing of the EEG lead field coefficient distribution.

**Figure S3.**
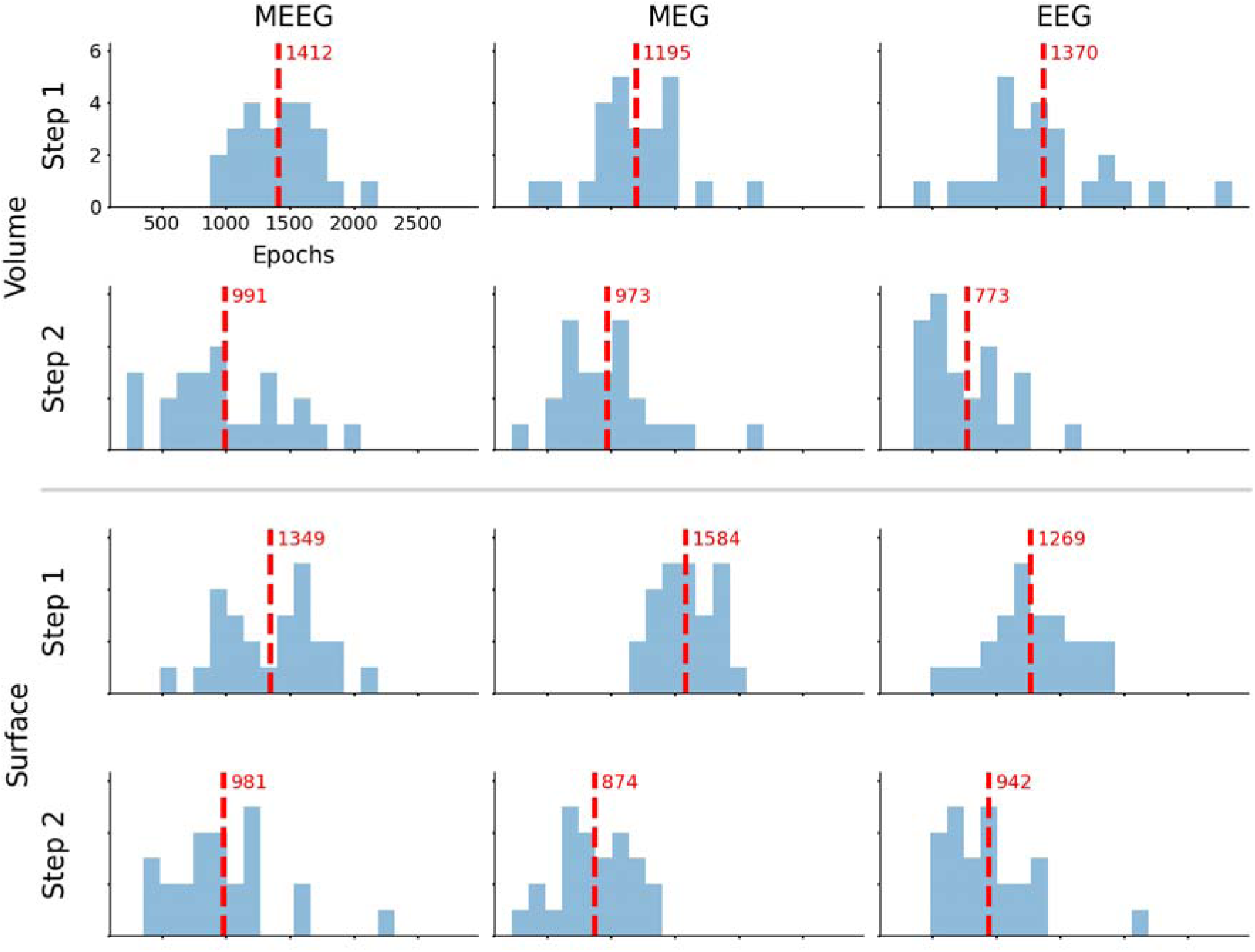
Histograms of the number of completed training epochs for RNNs without (step 1) and with L1 regularization (step 2) when fitted to MEEG, MEG-only and EEG-only. Vertical dashed lines and adjacent numbers indicate the mean number of epochs before stopping training. There were negligible differences among label types or between volume and surface source spaces. Transfer learning with L1 regularization applied to the source estimation layer (step 2) was completed in fewer epochs on average than pre-training the RNN model without additional constraints (step 1). Each histogram displays data from 25 models (five data scaling approaches times five seeds). All histograms are plotted with the same x-axis range.

**Table S1.**
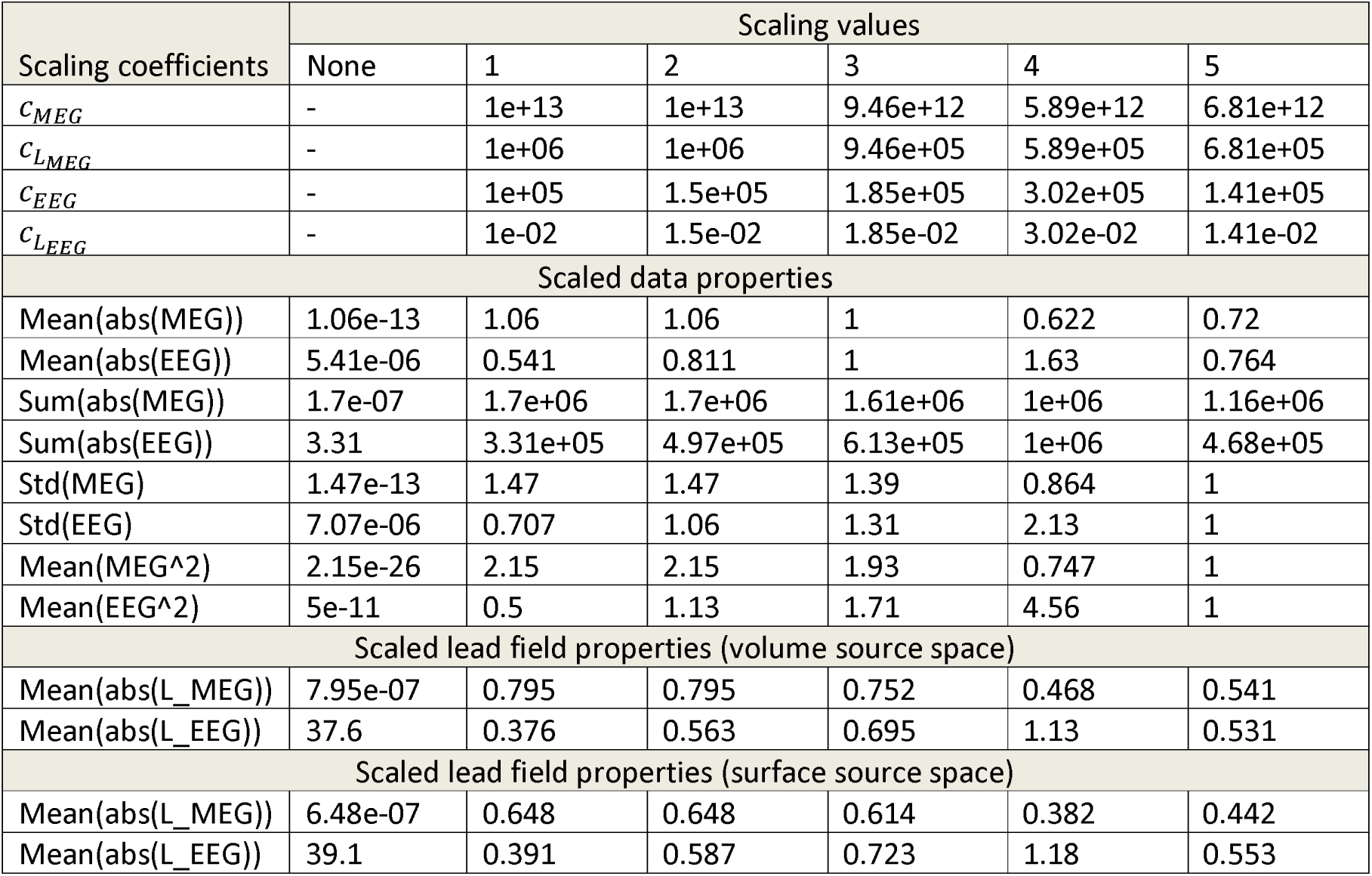
Summary of data and lead field scaling coefficients and corresponding scaled data and lead field properties.

**Figure S4.**
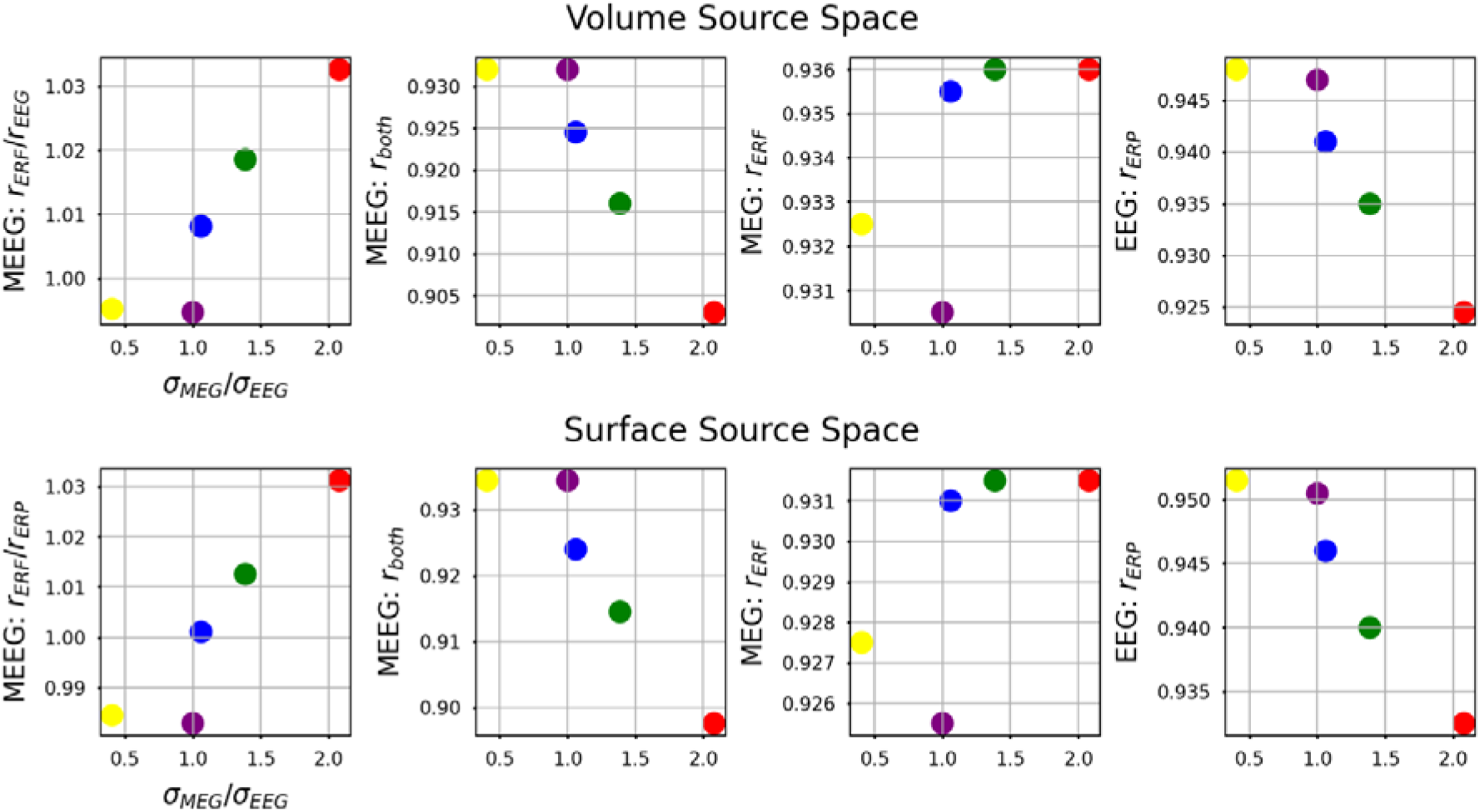
Correlation between reconstructed and ground-truth ERP and ERF waveforms from RNNs trained with different EEG and MEG scaling schemes. The ratio of standard deviations between scaled MEG and EEG data (σ_MEG_/σ_EEG_) used for training RNNs are plotted on the x-axes. Different scaling regimes are identified as follows; 1: red, 2: green, 3: blue, 4: yellow, 5: purple. The ratio of scaled MEG to EEG standard deviations was positively correlated with the ratio of correlation between reconstructed ERF to ERP waveforms from RNNs trained with MEEG.

**Figure S5.**
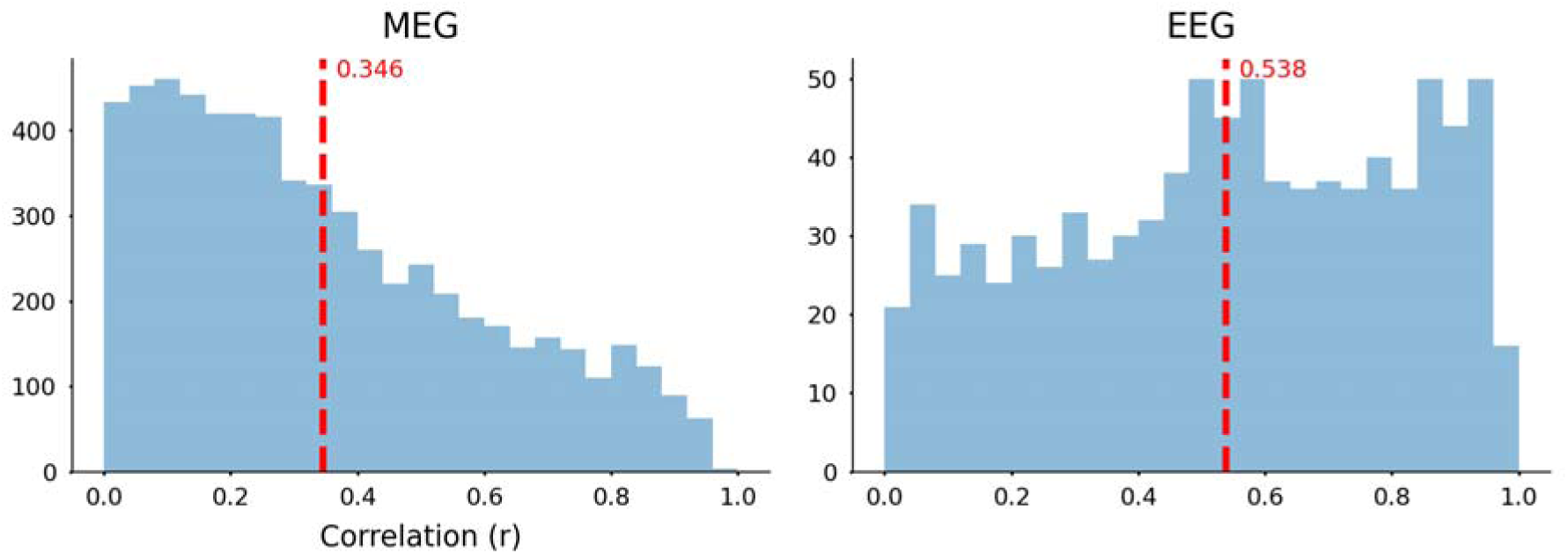
Distributions of correlation between MEG and EEG channels. EEG channels have higher average correlation among each other than MEG channels, reflecting differences in spatial spreading of magnetic and electric fields.

**Figure S6.**
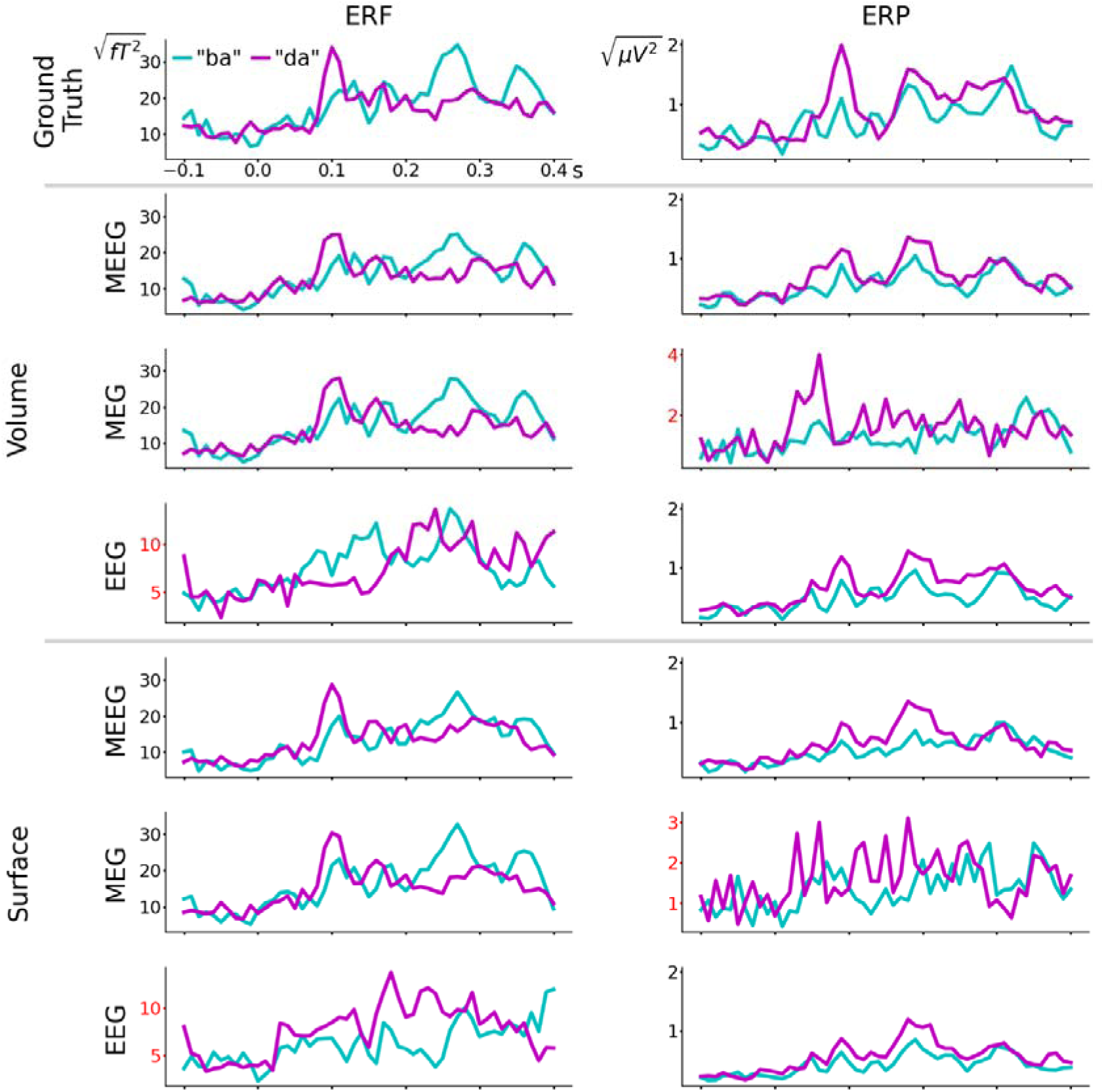
Reconstructed ERF and ERP waveforms from source signals estimated with eLORETA. Root-mean-squared amplitudes from all MEG and EEG channels are displayed in left and right columns, respectively. Most of the ERF plots on the left column are plotted with the same y-axis scale, although reconstructions derived from EEG data are plotted with different y-axis scales denoted by red labels. Conversely, ERP plots on the right column are plotted with a common y-axis scale unless derived from MEG data.

**Figure S7.**
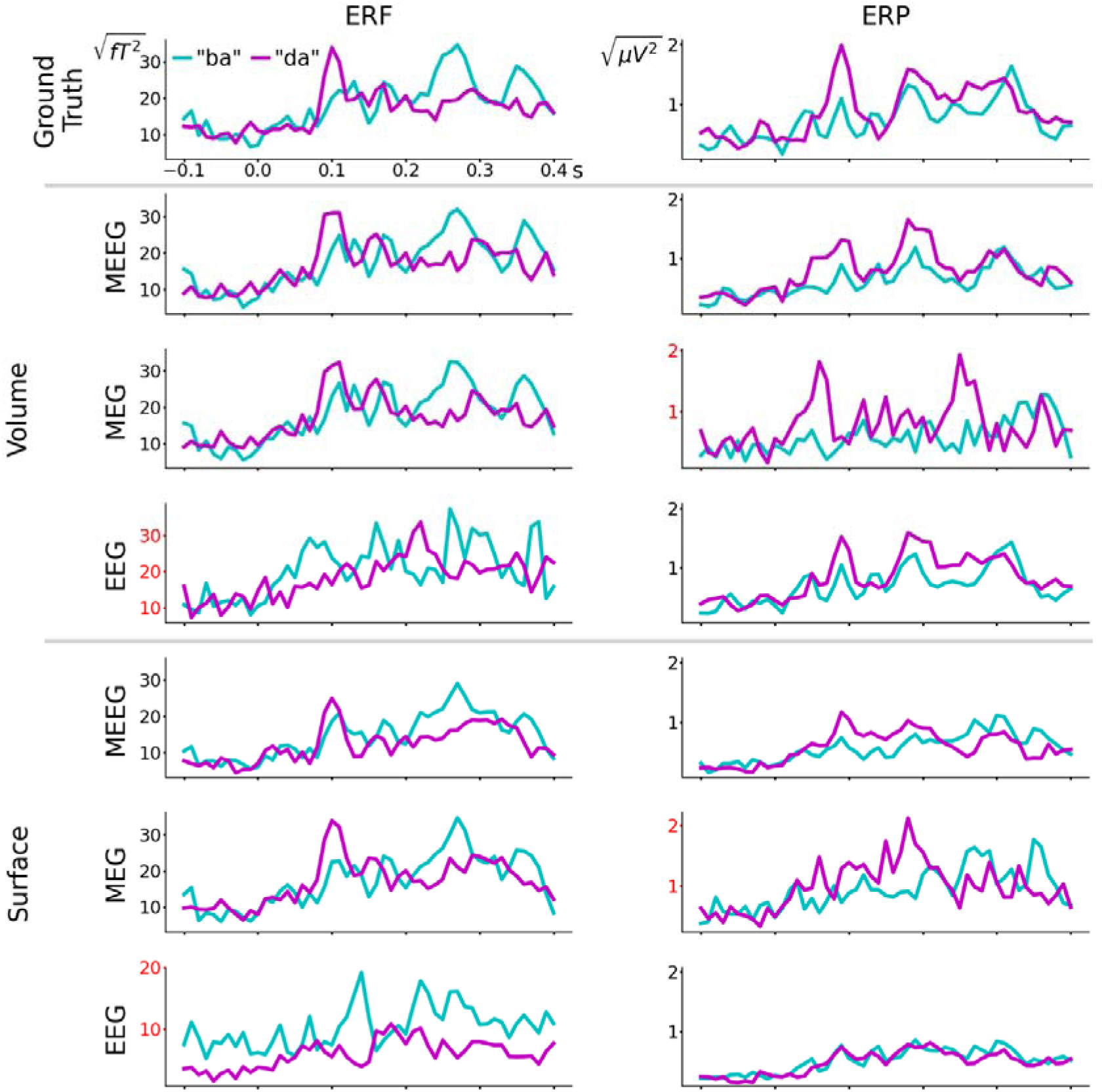
Reconstructed ERF and ERP waveforms from source signals estimated with MxNE. Root-mean-squared amplitudes are shown. Left column plots have a common y-axis scale if derived from MEEG or MEG data, but plotted on different y-axis scales derived from EEG data. Conversely, right column plots have a common y-axis scale if derived from MEEG or EEG data, but specific y-axis scales if derived from MEG data. Specific y-axis scales are highlighted with red labels.

**Figure S8.**
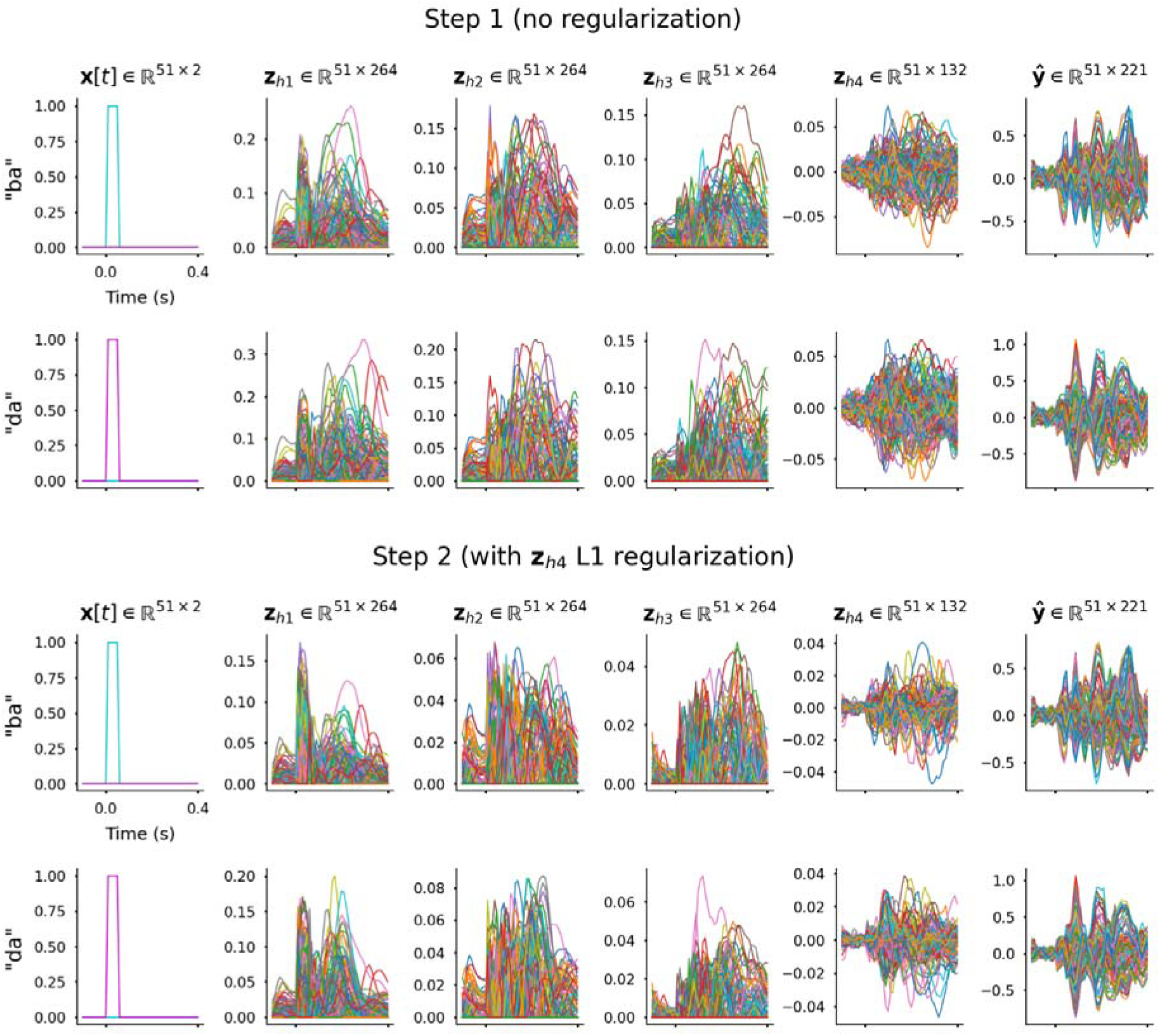
Example of RNN hidden unit activations. These signals were obtained from the best-fitting surface source space model in terms of summed ERF and ERP correlations, which was trained with data scaling regime 4, random seed 3. After training step 2, this model achieved r_ERF_ = 0.906 and r_ERP_ = 0.958. This depicts how simple representations of events are sequentially transformed into source signals that can account for observed ERF and ERP signals. Comparing upper and lower rows (step 1 vs. step 2), fine-tuning with L1 regularization applied to layer 4 activations (**z**_h4_) is seen to decrease hidden unit activations from layer 1 to layer 4. Y-axis units are activation value from individual artificial neurons in the RNN.

**Figure S9.**
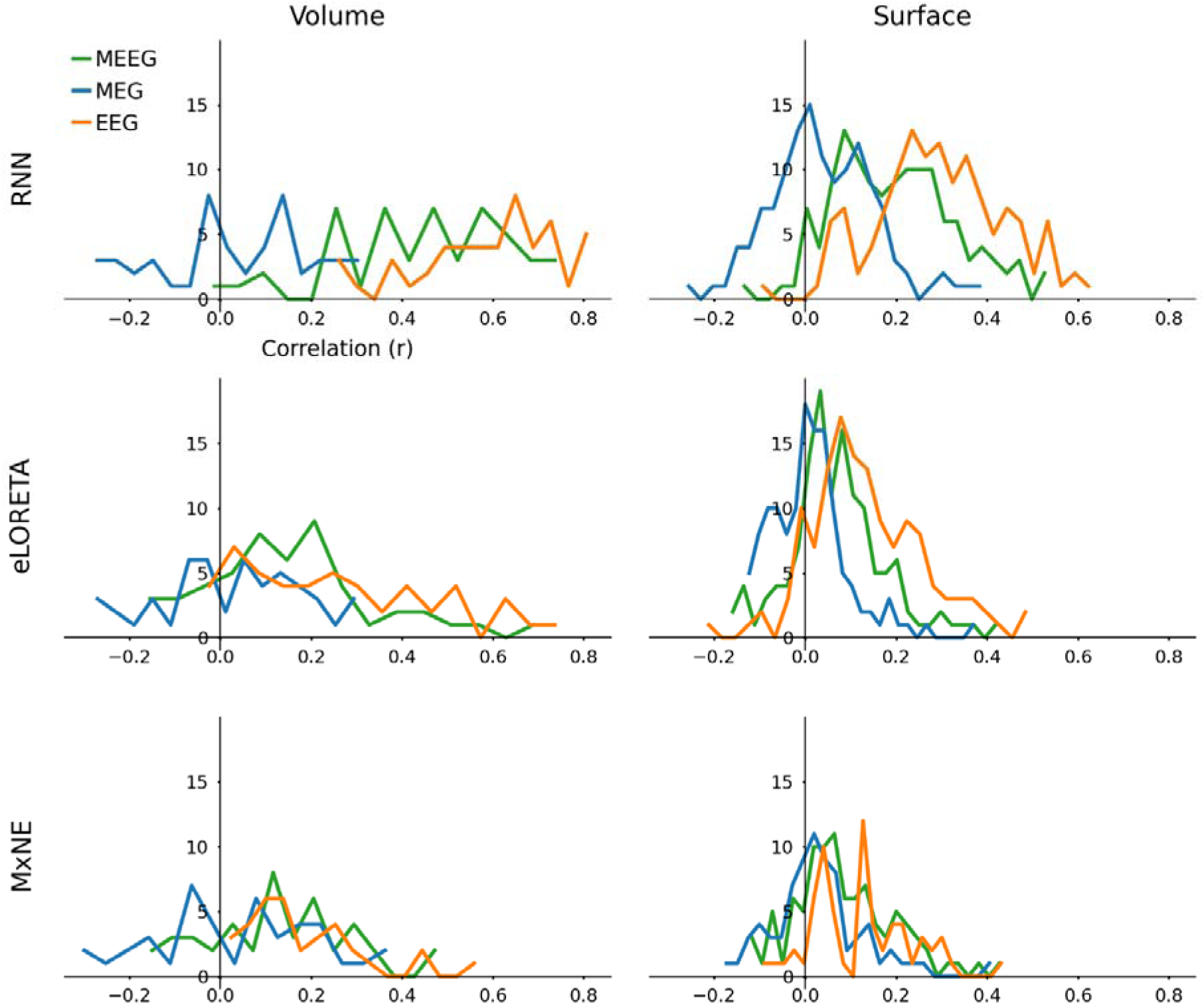
Distributions of correlation between ground truth and projected ERF and ERP waveforms. Projections were computed from individual estimated source signals and compared with both ERF and ERP waveforms for MEEG-derived sources, only ERFs for MEG-derived sources, and only ERPs for EEG-derived sources. Three direction components (x, y, and z) of volume sources (n=50) were projected simultaneously, whereas fixed-orientation surface sources (n=132) were projected individually.

**Figure S10.**
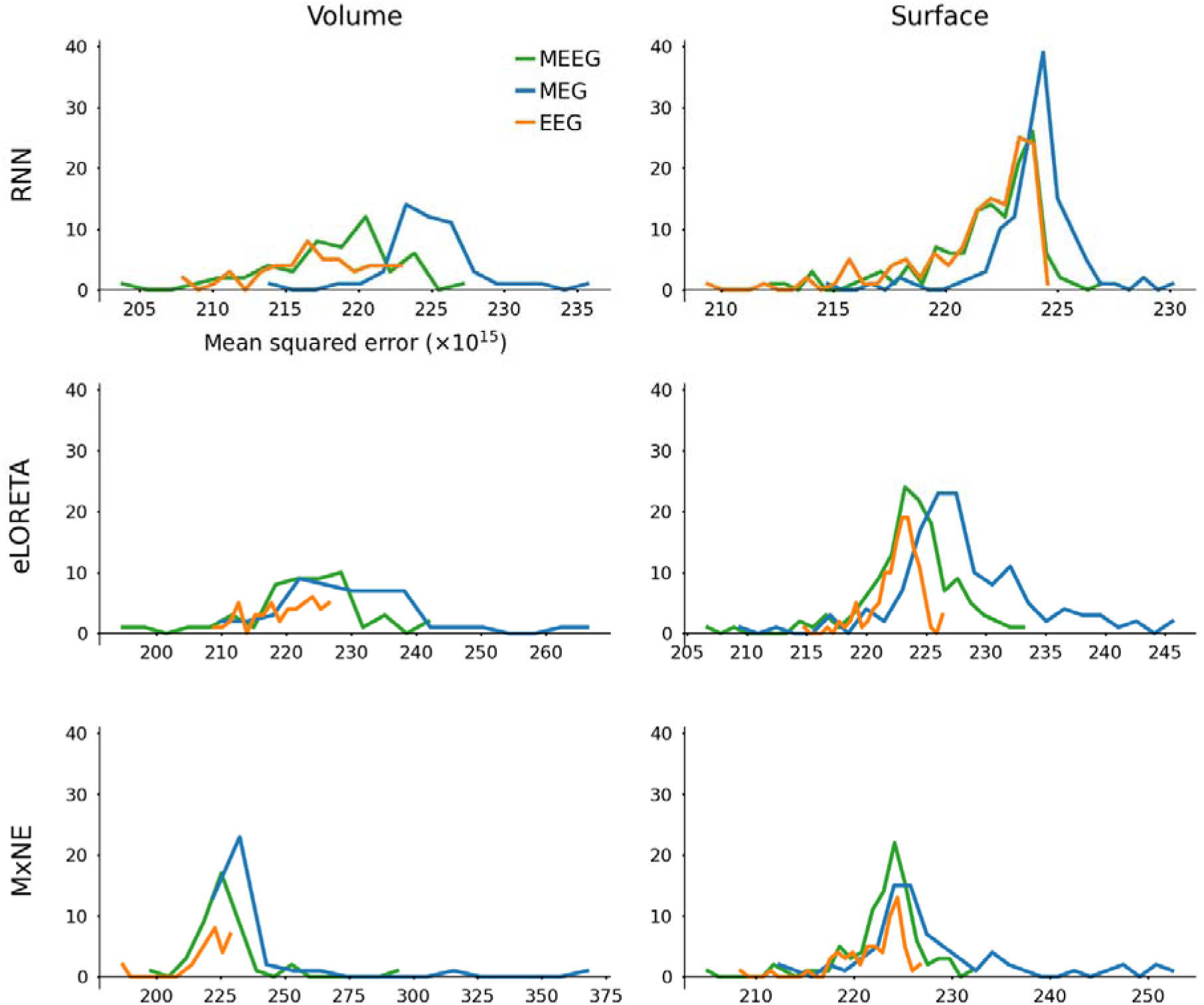
Distributions of MSE between ground truth and projected ERF/ERP waveforms. Plotted with different x-axes scales. Projections were made as described in the legend to Figure S8.

**Figure S11.**
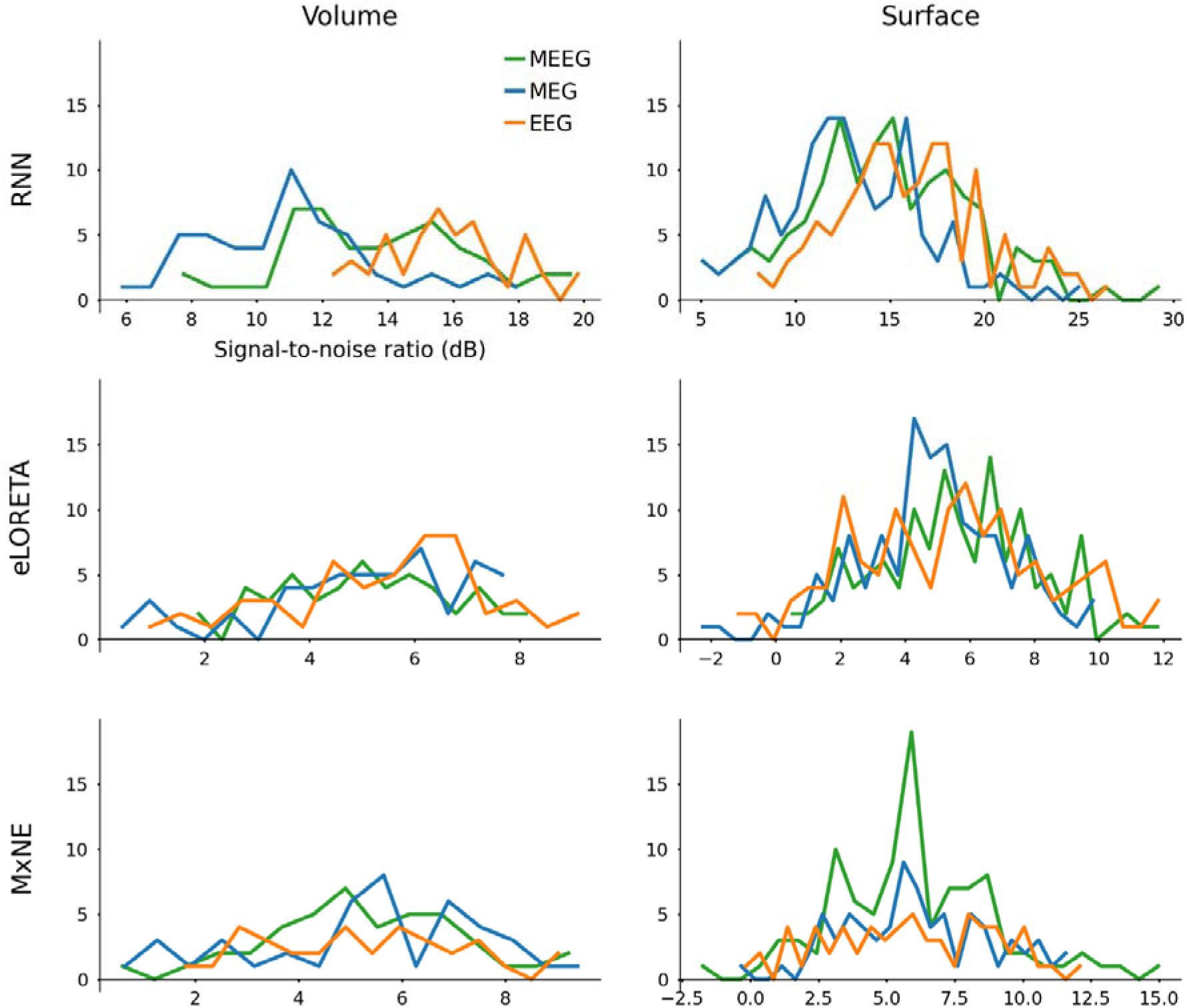
Distributions of SNR between ground truth and projected ERF/ERP waveforms. Plotted with different x-axes scales. Projections were made as described in the legend to Figure S8.

